# High Resolution Structural Insights into Heliorhodopsin Family

**DOI:** 10.1101/767665

**Authors:** K. Kovalev, D. Volkov, R. Astashkin, A. Alekseev, I. Gushchin, J. M. Haro-Moreno, A. Rogachev, T. Balandin, V. Borshchevskiy, A. Popov, G. Bourenkov, E. Bamberg, F. Rodriguez-Valera, G. Bueldt, V. Gordeliy

## Abstract

Rhodopsins are the most abundant light-harvesting proteins. A new family of rhodopsins, heliorhodopsins (HeRs), was recently discovered. In opposite to the known rhodopsins their N-termini face the cytoplasm. HeRs structure and function remain unknown. We present structures of two HeR-48C12 states at 1.5 Å showing its remarkable difference from all known rhodopsins. Its internal extracellular part is completely hydrophobic, while the cytoplasmic part comprises a cavity (’active site’), surrounded by charged amino acids and containing a cluster of water molecules, presumably being a primary proton acceptor from the Schiff base. At acidic pH a planar triangle molecule (acetate) is present in the ‘active site’ which demonstrated its ability to maintain such anions as carbonate or nitrate. Structure-based bioinformatic analysis identified 10 subfamilies of HeRs suggesting their diverse biological functions. The structures and available data suggest an enzymatic activity of HeR-48C12 subfamily and their possible involvement into fundamental redox biological processes.

## Introduction

Microbial and animal visual rhodopsins (classified into type 1 and 2 rhodopsins correspondingly) comprise an abundant family of seven transmembrane proteins that contain a covalently attached cofactor, the retinal^1–3^. Upon absorption of a photon, the retinal isomerizes triggering a series of conformational transformations correlating with functional and spectral states known as the photocycle^4, 5, 6^. Microbial rhodopsins are currently considered to be universal and the most abundant on the Earth light harvesting proteins. Before the year 2000, only rhodopsins from halophilic archaea had been known. About 30 years after the discovery of the first rhodopsin (bacteriorhodopsin, bR)^2^ metagenomics studies by Beja *et al.* led to the discovery in 2000 of a rhodopsin gene in marine Proteobacteria that was, accordingly, named proteorhodopsin (pR)^7^. After that seven thousand microbial rhodopsins were identified. They are present in all the three domains of life (bacteria, archaea and eukaryotes) as well as in giant viruses^4^. The discovery of channelrhodopsins^8^ led to the development of optogenetics, the revolutionary method for controlling cell behavior in vivo in which microbial rhodopsins play the key role^9–12^.

Several rhodopsins with new functions were discovered and characterized. Among the members of the rhodopsin family are light-driven proton, anion and cation pumps, light-gated anion and cation channels, and photoreceptors^5, 6, 13, 14^. Genomic and metagenomic studies dramatically expanded the world of rhodopsin sequences, some of which were found in unexpected organisms and habitats, for example, sodium-pumping rhodopsins (NaRs) in Flavobacteria^15, 16^. The widely spread presence and importance of pR-based phototrophy in the marine environment^17^ was identified. Recently, rhodopsins that function as inward proton pumps were discovered^18, 19^.

Despite diversity of their functions and differences in the structures, all these rhodopsins are oriented in the membranes in the same way. Their N termini always face the outside of the cells. Very recently Pushkarev *et al*. discovered a new large family of rhodopsins, named heliorhodopsins (HeRs), facing the cytoplasmic space of the cell with their N termini^20^. It was found that they are present in Archaea, Bacteria, Eukarya and viruses.

The function and structure of these heliorhodopsins are not yet known^21, 22^. Here we present high resolution crystallographic structures of a HeR (48C12), discovered in an actinobacterial fosmid from freshwater lake Kinneret^20^, corresponding to two states of the protein both solved at 1.5 Å resolution. The structures show an astonishingly large difference between the organization of the HeR and all known rhodopsins. For instance, the protein has a big cavity in the cytoplasmic part, containing the cluster of water molecules, which is likely to serve as proton acceptor from the retinal Schiff base (RSB). Despite such differences between HeRs and all the known rhodopsins the structure of 48C12 suggests that type 1 and HeR rhodopsins are evolutionarily related. Ten of 48C12 amino acids are highly conserved within all heliorhodopsins and we believe that its structure and the discussed mechanisms will be a basis for understanding the whole new abundant family and also the evolution of rhodopsins in general.

## Results

### Structure of 48C12 heliorhodopsin at neutral pH

48C12 was crystallized using the *in meso* approach similarly to our previous works^23^. Rhombic crystals appeared in two weeks and reached 150 μm in length and width with the maximum thickness of 20 μm. We have solved crystal structure of 48C12 heliorhodopsin at pH 8.8 at 1.5 Å. The crystals of P2_1_ symmetry contained two protein protomers organized in a stable dimer per asymmetric unit (Fig. S8). The high-resolution structure reveals 233 water molecules and 31 lipid fragments.

The protein has an architecture which differs from that of known rhodopsins. Although similarly to type I rhodopsins, each 48C12 protomer has seven transmembrane α-helices, connected by three extracellular and three intracellular loops, some of the loops are relatively large and have certain secondary structure (Fig. 1). Particularly, the extracellular AB-loop of 48C12 (residues 34-64) is ∼40 Å long and forms a β-sheet with the length of ∼17 Å (Fig. 1B). It extends in the direction of the second protomer of the dimer remaining parallel to the membrane surface and thus covers the extracellular surface of the nearby molecule (Fig. 1, Fig. S1). The intracellular BC-loop comprises 14 residues (86-98) and forms an α-helix with the length of ∼18 Å (Fig. 1C). Other loops and N- and C-termini although not forming regular secondary structures, are well-ordered and therefore are completely resolved.

**Fig. 1.**
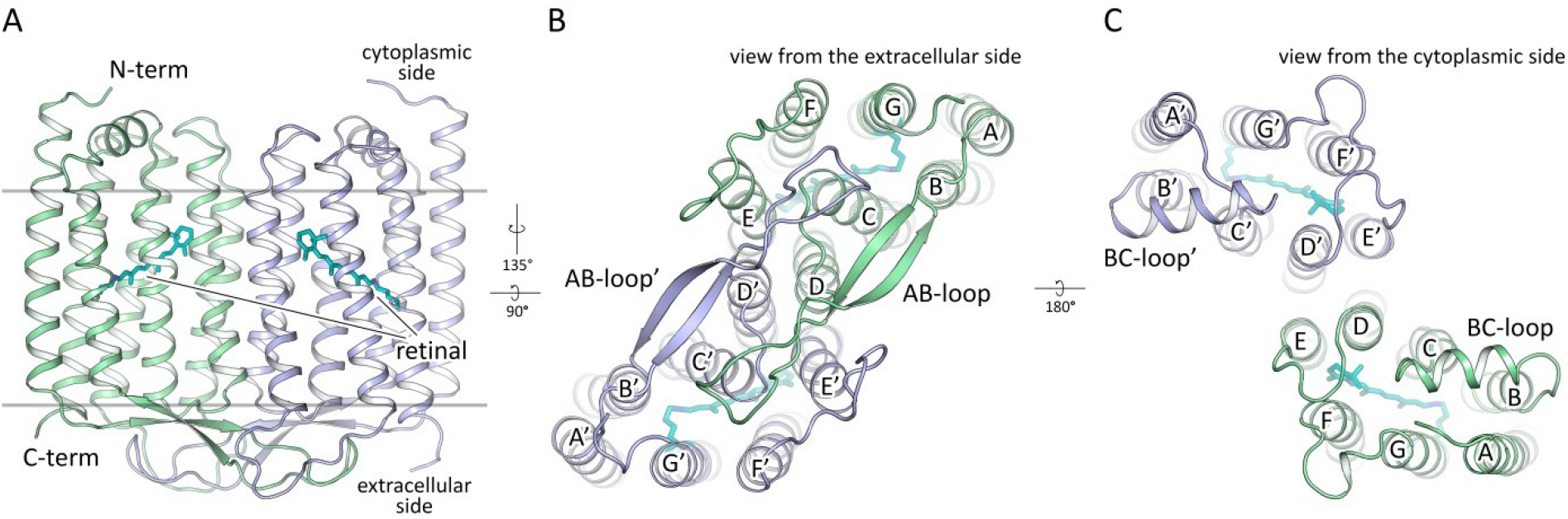
Overall architecture of 48C12 dimer. **A.** Side view of the dimer. Hydrophobic/hydrophilic membrane boundaries are shown with gray lines. **B.** View from the extracellular side. **C.** View from the cytoplasmic side. Cofactor retinal is colored teal.

### Dimeric structure of the protein

48C12 protomers in a dimer interact via helices D and E (Fig. 1B, C; Fig. S3), with a broad hydrophobic interface in the middle part (inside the membrane) and interactions between polar residues, specifically Asp127 and Tyr179’ at the extracellular and Tyr151 and Asp158’ at the cytoplasmic sides of the membrane. Tyr179’ side chain is additionally connected through a hydrogen bond to the main chain of the AB-loop of the neighbor protomer (nitrogen of Thr44). The AB-loop itself almost does not interact directly with the neighbor protomer, although it is stabilized by several hydrogen bonds mediated by numerous water molecules located on the extracellular surface of the dimer.

Several well-ordered lipid molecules are present in the structure, surrounding the protein dimer (Fig. S4). Two of them permeate inside nearby protomer between helices E and F near the β-ionone-ring of the retinal cofactor with the hydrocarbon tails. Surprisingly, the pocket of the hydrocarbon chain comprises polar amino acids Asn207, Asn138 and one water molecule. Asn207 is also exposed to the surface of the extracellular part of the 48C12 protomer in the middle of membrane and is highly conserved within HeRs.

The protomers within the 48C12 dimer are similar (RMSD between protomers 0.144 Å), however differs in the EF-loop organization, α-helical BC-loop location and the 3 Å displacement of the cytoplasmic end of helix A (Fig. S7). Consequently, positions of several residues inside the protomers are slightly varied. Since general features of the heliorhodopsin structure are the same in both molecules, we will describe mostly the protomer A. Nevertheless, we will also describe the differences between the protomers where appropriate.

### Mechanism of topological inversion of heliorhodopsins

It is known that insertion and folding of membrane proteins is guided by the “positive-inside rule”^24^. Using the structure of 48C12 heliorhodopsin, we analyzed the location of positively and negatively charged residues in the cytoplasmic and extracellular domains of the protein and compared it to bacteriorhodopsin (bR) (Fig. 2). Notably, in 48C12 all the positively charged residues are located exclusively at the cytoplasmic side of the protein, which is consistent with the “positive-inside rule”^24^. Importantly, some of these residues, such as Arg91, Lys218, Lys222, and Arg231 are highly conserved in the subfamily of 48C12 (Fig. S14). In addition, unlike bR, the HeR contains only negative amino acids in the extracellular polar part of the proteins, which is characteristic for this subfamily. Thus, we suggest that HeRs follow the “positive-inside and negative-outside rule” rather than just the “positive-inside rule”.

**Fig. 2.**
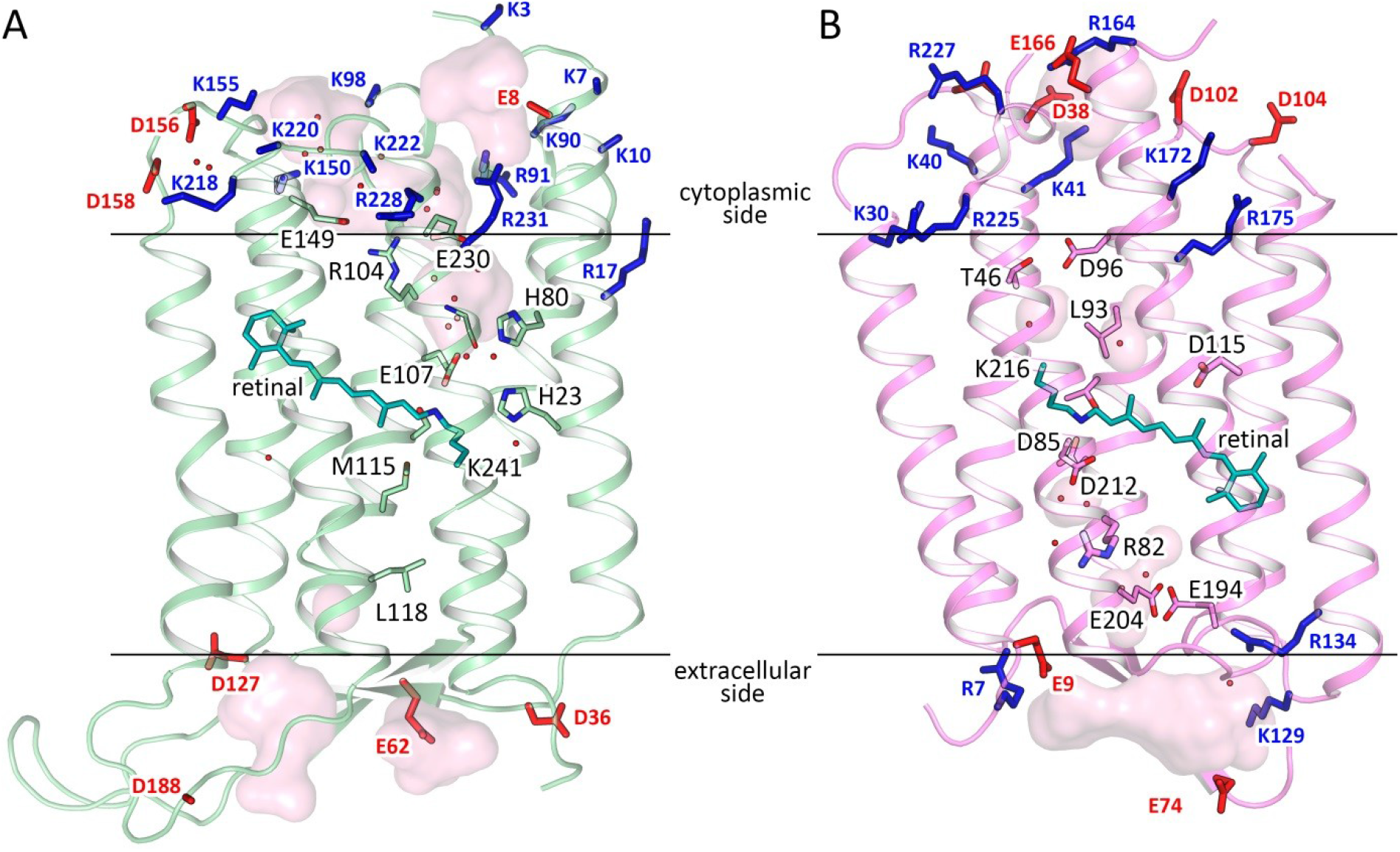
Comparison of 48C12 (green) and bR (purple, PDB ID 1C3W). **A.** Side view of the 48C12 heliorhodopsin. N-term is at the cytoplasmic side of the membrane. **B.** Side view of the bR. N-term is at the extracellular side of the membrane. Hydrophobic/hydrophilic membrane boundaries are shown with black lines. Positively and negatively charged residues on the protein cytoplasmic and extracellular surface are shown in blue and red, respectively

### Structure of the extracellular region

As heliorhodopsins are topologically inverted in the membrane relative to type I rhodopsins, the extracellular part of 48C12 corresponds to the cytoplasmic part of classical microbial rhodopsins, such as bR. However, in 48C12 the internal region, embedded in the extracellular half of lipid bilayer, is completely hydrophobic and compact and does not comprise any charged or polar amino acids and solvent-accessible cavities (Fig. 3C, Fig. 4). Hereafter, we denote this part as the hydrophobic extracellular region. Nevertheless, several clusters of polar amino acids are located at the extracellular half of the protein inside the membrane, but on the outer surface of the protein. Helices A and G interact by hydrogen bonding of Gln26 with Ser242 and Trp246, while helices F and G are also connected by a hydrogen bond between Gln247 and Ser201. We suggest that these interactions keep in the internal hydrophobic configuration at the extracellular side. The absence of any charged or/and polar amino acids inside the region may explain the absence of any proton/ion pumping by 48C12^20^.

**Fig. 3.**
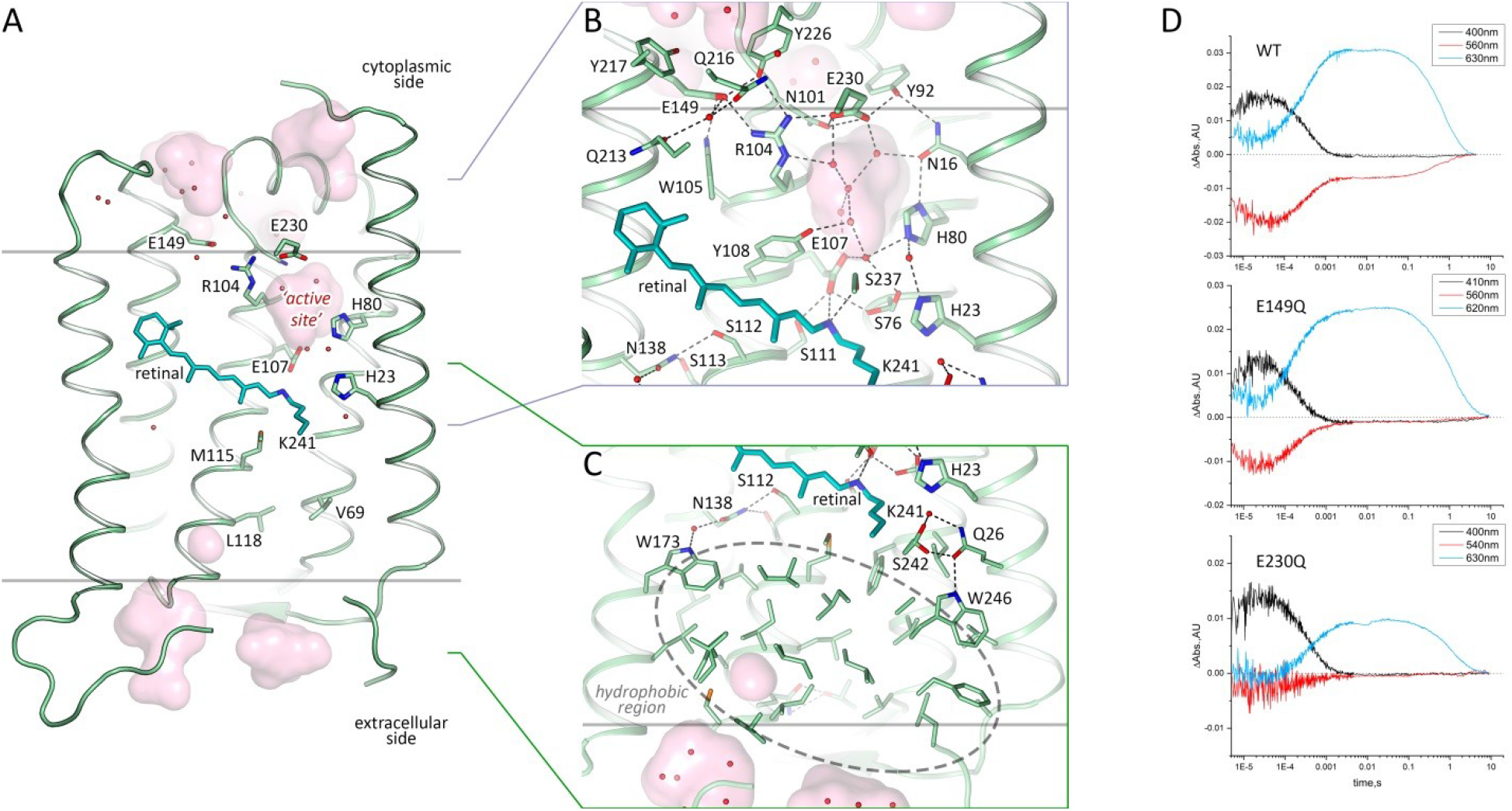
Structure of 48C12 protomer. **A.** Side view of the protomer in the membrane. **B.** Detail view of the cytoplasmic part. **C.** Detail view of the extracellular side and hydrophobic region. Cofactor retinal is colored teal. Hydrophobic/hydrophilic membrane boundaries are shown with gray lines. Cavities are calculated with HOLLOW and shown in pink. Charged residues in 48C12 are shown with thicker sticks. Helices F and G are not shown. D. Time evolution of transient absorption change of photo-excited 48C12 wild type (WT) and E230Q and E149Q mutant forms.

**Fig. 4.**
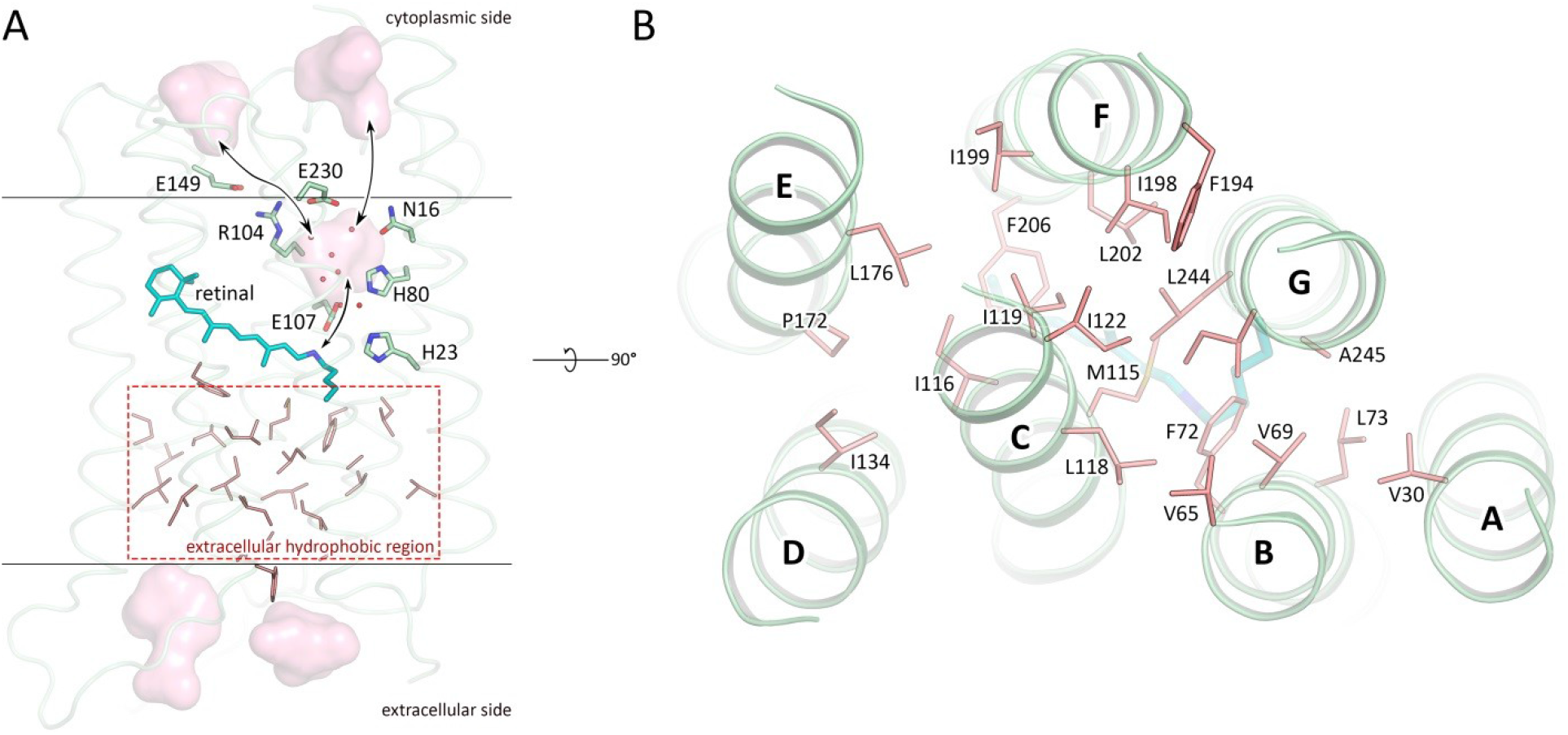
Hydrophobic residues in the extracellular part of 48C12. **A.** Side view of the 48C12 protomer. Residues, comprising the extracellular hydrophobic region are colored red. The region is embedded in the extracellular half of the lipid bilayer and is contoured with dashed red rectangle. Membrane core boundaries are shown with black lines. Black arrows indicate putative ways of the connection between inner cavity (‘active site’) and cytoplasmic side and RSB. Cavities are colored pink. Water molecules in the inner cavity are shown with red spheres. **B.** View on the hydrophobic region from the extracellular surface of the protein. Loops are hidden for clarity. Hydrophobic residues in the extracellular internal part of the 48C12 protomer are colored red. Cofactor retinal is colored teal.

### Retinal binding pocket and cavity in the retinal Schiff base region

The retinal binding pocket of 48C12 (Fig. 1 and 2) is also different from that of microbial rhodopsins with known structures. Near the retinal molecule helices C and D are connected by hydrogen bonding of Asn138 (analog of Asp115 in bR and Asp156 in ChR2) with Ser112 (analog of Thr90 in BR and Thr128 in ChR2) and Ser113. The Asn138 side chain is also stabilized by hydrogen bonding with Trp173 through a well-ordered water molecule (Fig. 3C). In the region of the β-ionone-ring of the retinal molecule only two residues (Met141 and Ile142) are similar to those in bR (Fig. S2). Although many of the residues of the pocket walls remain aromatic in 48C12, there are notable alterations such as for example Phe206 in the position of Trp182 in bR, Trp105 instead of Tyr83, Tyr108 in the place of Trp86. All these residues are highly conserved in heliorhodopsins. Interestingly, polar Gln213 (in the position of Trp189 in bR) is located close to the β-ionone-ring.

The Schiff base is surrounded by an unusual, for the rhodopsins of type I, set of residues: for example, Ser237 replaces Asp212 (extremely conserved aspartate in type I rhodopsins), Glu107 replaces Asp85, His23 replaces Met20, the bulky Phe72 replaces Val49, Met115 replaces Leu93, Ser76 replaces Ala53. In this configuration, RSB is hydrogen bonded directly to Glu107 (RSB counter ion) and Ser237. The Glu107 sidechain is stabilized by two serine residues (Ser76 and Ser111).

The distinctive feature of 48C12 heliorhodopsin is the presence of a large hydrophilic cavity in the vicinity of the Schiff base between residues Glu107 and Arg104 (analog of Arg82 in bR). The cavity is separated from the cytoplasmic bulk with the only side chain of Asn101 (Fig. 3) and is surrounded by polar residues Glu107, His23, His80, Ser237, Glu230, Tyr92, Asn16, Asn101, Tyr108 and filled with 6 water molecules (Fig. 3B). The listed amino acids together with the water molecules create a dense hydrogen bonding network, which protrudes from the RSB to Arg104. The Arg104 side chain is pointed towards the cytoplasm and is stabilized by Glu230, Glu149 and also Tyr226. It should be noted, that in protomer B there is an alternative conformation of Arg104, Glu230 and Tyr226, which, however, does not affect the shape of the cavity. The Glu149 side chain is additionally stabilized by Trp105 and by a water-mediated hydrogen bond to Gln216 and Gln213. The calculation of the hydrophilic/hydrophobic membrane boundaries shows that Glu149 is located out of the hydrophobic part of the membrane (Fig. 3A) and can be accessed from the cytoplasmic bulk, which is also proved by the cavities calculations using HOLLOW^25^. In protomer B the accessibility of Glu149 from the bulk is lower, mostly because of slight alterations of the helices positions. Importantly, all the residues mentioned in this paragraph are highly conserved within all the known heliorhodopsins (Fig. S11, S13, S15). This fact together with their structural roles points towards their functional importance.

As it was shown in previous studies, His23, His80 and Glu107 do not act as a proton acceptor from the RSB, however His23 and His80 are important for proton transfer ^20, 21^. Our structure shows that the also charged amino acids E149 and E230 are connected via a continuous network of hydrogen bonds to the RSB, but have not yet been studied. To understand better their roles for the heliorhodopsin functioning we produced E149Q and E230Q mutants and studied the properties of their photocycles. First of all, we should stress that the mutants are not stable under purification. Moreover, E230Q degrades quickly upon illumination even being in the lipid membranes. It indicates that both amino acids are important for the protein stabilization. We measured the transient absorption of the mutants with the solubilized (not purified) protein (Fig. 3D). The M formation was observed for both mutants, therefore neither Glu149 nor Glu230 are the proton acceptor. However, the O-state decay in mutants is more than twice longer than that of wild type protein (Fig. 3D), which indicated the involvement of Glu149 and Glu230 in the protein work. Thus, none of charged amino acids, surrounding the inner cavity at the cytoplasmic part of protein is a proton acceptor. Taking into account all facts (also the absence of charged amino acids in the hydrophobic extracellular internal part of the protein and that the proton is not transiently released to the aqueous phase^20^ we conclude that the only candidate for the proton acceptor is water cluster in the cavity. Indeed, water molecules were shown to play key role in functioning of microbial rhodopsins^26^. Thus, we suggest that proton is stored in the aqueous phase of the cavity after its release from the RSB and is returned to the RSB in the end of 48C12 photocycle. Hereafter, we denote the internal cavity at the cytoplasmic part of the 48C12 as an ‘active site’.

### Structure of 48C12 at acidic pH

While the biological function of HeRs remains unknown, thorough study of different 48C12 states is of great benefit. To investigate the conformational rearrangements in the heliorhodopsin associated with pH decrease, we also solved the crystal structure of 48C12 at 1.5 Å using the crystals grown at pH 4.3. Indeed, pH of the surrounding solution affects the functionality and structure of microbial rhodopsins due to protonation or deprotonation of the key residues ^27–29^. Moreover, it was shown for BR that the structure of the protein at acidic pH is similar to that of its M intermediate state ^30^. Thus, analysis of heliorhodopsin structure at low pH may be of high importance for understanding of its biological function and possible rearrangements in protein structure during photocycle. While the crystal packing is the same as in crystals grown at neutral pH with one 48C12 dimer in the asymmetric unit, the crystals were colored blue (maximum absorption wavelength of 568 nm) at acidic pH, while at neutral pH they were violet (maximum absorption wavelength of 552 nm), which corresponds to the color of the wild-type protein in solution under the same conditions^21^ (Fig. S8). We designate these two 48C12 forms as blue and violet, respectively. The color shift is presumably caused by the protonation of the Glu107 residue^20^, thus we suggest that Glu107 is protonated in the blue form. Key differences between two structures are shown in Fig. 5. In general, the backbone organization is the same at both acidic and neutral pH values (RMSD between models 0.158 Å), however the cytoplasmic parts of helices A and B are displaced for 1 to 2 Å respectively in the blue form (Fig. S7).

**Fig. 5.**
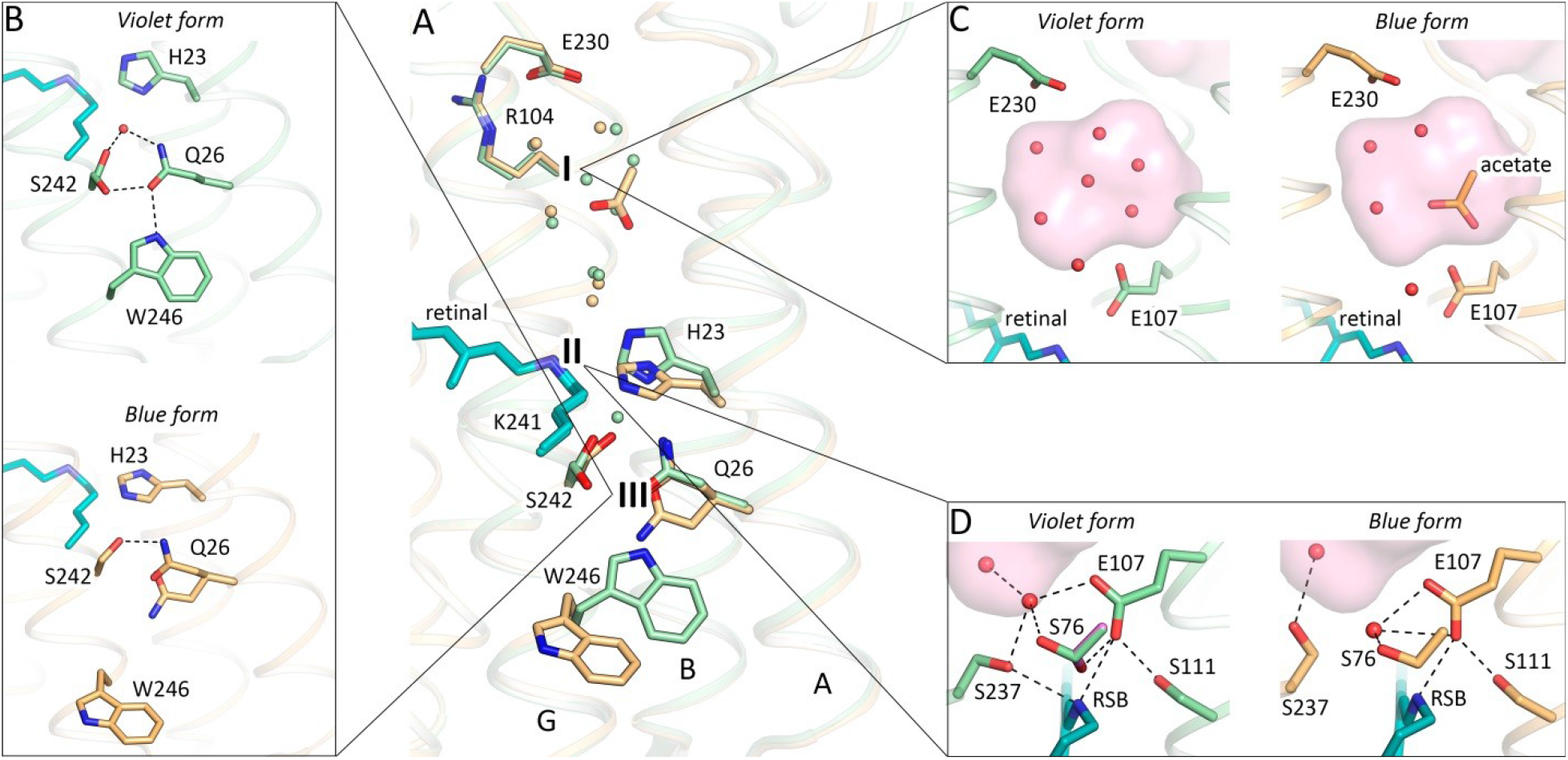
Comparison of the violet (green) and blue (orange) forms of 48C12. **A.** Alignment of two models. Three most notable differences between two structures are present: I –the cavity at the cytoplasmic side; II – rearrangements of residues near the RSB; III – loss of the water molecule between His23 and Ser242 in the blue form and rearrangements of the Gln26 and Trp246 side chains. Water molecules are shown with the spheres and colored green and orange, corresponding to the violet and blue forms of 48C12, respectively. **B.** Detailed view of the Ser242-Gln26-Trp246 cluster and His23 in the violet and blue forms. **C.** Detailed view of the cavity (active site) at the cytoplasmic side in the violet and blue forms. **D.** Detailed view of the RSB and surrounding residues in the violet and blue forms. In violet form Ser76 shapes two alternative conformations (second is colored magenta for clarity). Cavities are colored pink.

At the cytoplasmic side the main difference is in the organization of water molecules inside the big cavity near the RSB (Fig. 5C). The hydrogen bonds network propagating from the RSB to Arg104 and Glu230 is present in both models. Interestingly, the difference F_o_-F_c_ electron densities at 1.5 Å resolution indicate the presence of a triangle molecule in the ‘active site’ (Fig. S9F). As the crystallization buffer contained only one molecule of triangle geometry, acetate anion, the densities were fitted with an acetate (CH_3_COO^-^) molecule (Fig. 5, Fig. S9E). It fits the density, however, the acetate can mediate only two hydrogen bonds instead of three bonds necessary to fit the environment of two water molecules 3 and 5 and Glu107 (Fig. 5C, Fig. S9C, D). Other molecules that could fit the triangle density and create three hydrogen bonds with water molecule 3 and 5 and Glu107 could be nitric acid (HNO_3_^-^) or bicarbonate (HCO_3_^-^) (Fig. S9C, D). This fact indicates the ability of 48C12 heliorhodopsin to bind anions of triangle geometry inside the ‘active site’.

While at neutral pH RSB is stabilized through hydrogen bonding to Ser237 and Glu107, at acidic pH it is still bound to Glu107, however is slightly shifted towards Ser111, thus weakening connection to Ser237 (Fig. 5D). Ser76 is in a single conformation at acidic pH and does not stabilize Glu107 anymore. Ser237 flips from the RSB towards the cavity at the cytoplasmic part of the protein (Fig. 5D). At the same time His23 is reoriented compared to the structure at neutral pH, and forms a hydrogen bond with Ser76 in the blue form. The reorientation of His23 may be caused either by protonation of Glu107^20^ or by protonation of His23 itself, or the combination of the both. Nevertheless, the reorientation of His23 towards the extracellular side results in loss in the blue form of HeRs of the water molecule, which is coordinated by Gln26 in the purple form (Fig. 2C). The organization of the Ser242-Gln26-Trp246 cluster, located at the extracellular half of the protein surface inside the hydrophobic membrane, is also disturbed (Fig. 5B). Prominently, Trp246, conserved in most HeRs (Figs. S11, S13, S14) and exposed to the surrounding lipid bilayer, loses the hydrogen bond to Gln26 and reorients in the blue form of 48C12. Such reorientation might trigger a signal transduction cascade, if heliorhodopsins are light sensors^20^, similarly to sensory rhodopsins II, where there is also an aromatic amino acid in the helix G, Tyr199, which controls the signal transducer protein^31^. Alternatively, a latch-like motion of Trp246 might create a defect in the surrounding lipid membrane and open a pathway towards Gln26, His23 and the retinal.

### Structure-based bioinformatic analysis of heliorhodopsins

The structures of 48C12 allowed us to identify amino acid residues, comprising the key regions of the heliorhodopsin (Fig. 6). Based on the comparison of these amino acids in different HeRs, we classified heliorhodopsins in ten subfamilies with potentially different properties. The subfamilies are presented in a phylogenetic tree (Fig. S11). The groups which contain less than 10 members were merged into “Unsorted proteins”.

**Fig. 6.**
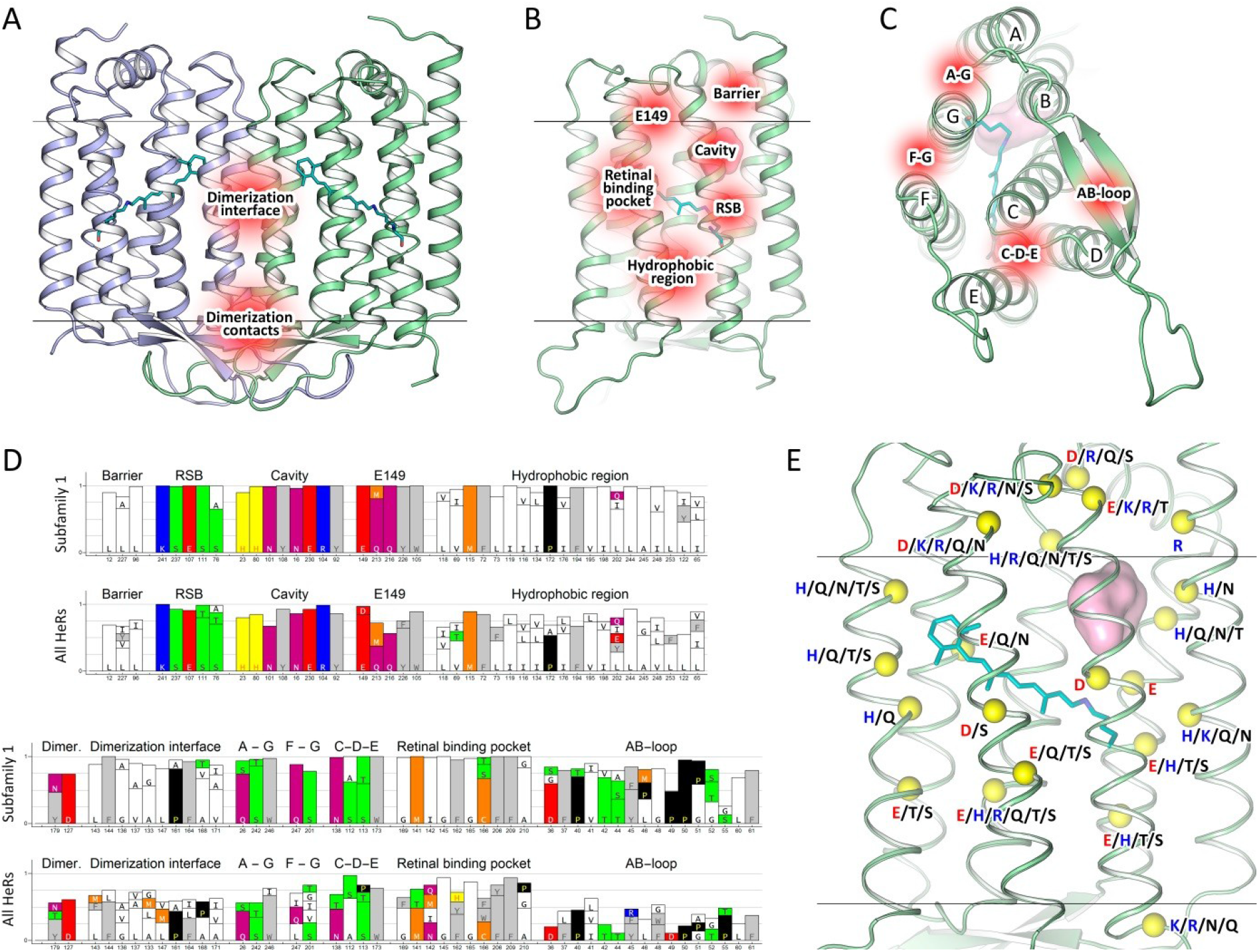
Key regions of 48C12 and heliorhodopsins family identified by structure-based bioinformatical analysis. **A.** View of the 48C12 dimer with identified regions (shown in red) comprised of conservative residues of the dimerization interface (polar, responsible for contacts between protomers and hydrophobic, responsible for hydrophobic interaction inside the membrane) **B.** View of the 48C12 protomer with the key regions (shown in red), comprised of conservative residues. **C.** View of the 48C12 protomer with the polar clusters (shown in red), shown to be conservative among subfamily 1 and most of other subfamilies. **D.** Most conservative residues among subfamily 1 and all HeRs, comprising the key regions of 48C12. **E.** The location of the residues in 48C12, which are probable analogues of charged or polar residues in other subfamilies of HeRs, were selected using a sequence alignment (Fig. S16, S17). The backbone carbon atoms of these residues are shown with yellow spheres. The polar cavity in the cytoplasmic part is shown with a pink surface. Cofactor retinal is colored teal.

*The group of 48C12 (subfamily 1)* is the largest and comprises 195 proteins from all 479 unique sequences of HeRs currently available^20, 32^. The majority of HeRs of subfamily 1 have bacterial origin, with most of them from *Actinobacteria*. However, representatives of the subfamily are also found in *Chloroflexi* and *Firmicutes* of the *Terrabacteria* group and also in *Proteobacteria* and the PVC group. The host of the unique protein A0A0L0D8K8 is eukaryotic *Thecamonas trahens*. Importantly, the sequences belong to both gram-positive and gram-negative bacteria, which is inconsistent to previously made conclusion^22^.

Those residues that are conservative in the most of the proteins were identified (Fig. S14). The alignment of 14 most distinct heliorhodopsins of subfamily 1 is shown in Fig. S13. Using the structure of 48C12 as reference we identified the following regions of protein comprised of conserved residues as potentially important for the function of 48C12 and correspondingly for the whole 48C12 subfamily and in some cases for all HeRs (Fig. 6).

Namely, heliorhodopsins have a conservative pattern of the residues that stabilize the RSB (Ser237, Glu107, Ser111 and Ser76) (Fig. S15, S16). The big inner cavity in the cytoplasmic part and surrounding it the charged and polar residues (His23, His80, Asn101, Tyr108, Asn16, Glu230, Arg104, Tyr92) together with residues Leu12, Leu96, and Leu227 forming a hydrophobic barrier between the cavity and the cytoplasm are almost completely conserved in subfamily 1 (Fig. S15, S16). The polar region near Glu149 and Arg104 is also conservative (Glu149, Gln216, Tyr226, Trp105, Gln213) (Fig. S15, S16). We found that the common feature of the heliorhodopsins is the hydrophobic organization of the extracellular internal part (Fig. 3, Fig. S15, S16). Indeed, only a few heliorhodopsin subfamilies have members with charged or polar residues in the region. This fact is very interesting from the functional point of view and will be discussed in the latter paragraphs. As it was already mentioned, 48C12 has three clusters comprised of polar residues (Gln26/Ser242/Trp246, Gln247/Ser201 and Ser112/Ser113/Asn138/Trp173), which are structurally important presumably for the helices interactions and stabilization of the protein. Importantly, many residues of the characteristic for 48C12 long AB-loop and dimerization interface are conserved within subfamily 1. To identify whether the same regions and residues are conserved within other HeR subfamilies, we performed additional bioinformatic analysis of the whole family.

*The most conservative residues in HeR family* are similar to those of subfamiliy 1 with some variations. The filling of the cytoplasmic part, particularly the RSB region, the inner cavity in the cytoplasmic part, the hydrophobic barrier separating the cavity from the cytoplasm, the region near Glu149, as well as the hydrophobic extracellular configuration are highly conserved within all HeRs (Fig. S14, S16). These regions include such polar and charged residues as Ser237, Glu107, Ser111, Ser76, His23, His80, Asn101, Tyr108, Asn16, Glu230, Arg104, Tyr92, Glu149, Gln216, Tyr226, Trp105 and Gln213, which were shown to be structurally important in 48C12 (Fig. S14, S16). Indeed, only a few heliorhodopsin subfamilies have variations in the listed parts (Fig. S16, S17). It should be noted, that although Gln213 is almost completely conserved among heliorhodopsins of subfamily 1, methionine is an often variant for this position in heliorhodopsins. In addition, the analogues of the residues, comprising the clusters at the surface of 48C12 (Gln26/Ser242/Trp246 and Gln247/Ser201) are present in most of the HeRs.

*The differences between heliorhodopsin subfamilies* were identified by a comparison of the residues, comprising structurally important regions in 48C12. In general, amino acids responsible for dimerization are not conserved in all HeRs. However, in most cases analogues of Tyr179 and Asp127 are present (except subfamily 2) but hydrophobic residues of the dimerization interface are different in almost all the groups. The AB-loop is conserved only within some subfamilies, but varies notably from group to group in size and amino acid composition. Despite this, residue Pro40 is highly conserved among all HeRs and is part of a β-sheet of the AB-loop of 48C12.

*Subfamily 2* comprises 19 members and mostly consists of viral proteins, but there are two representatives of *Euryarchaeota*; the bacterial *PVC* group and eukaryota are also presented with 1 protein. We found that this group is the most distinct from all others especially in the organization of the extracellular part and the retinal binding pocket. Interestingly, one of the members of this group has two Asn residues near the cytoplasmic inner cavity in the positions of His23 and His80 of 48C12. A lot of its members have glutamate in helix F in the position of Leu202 in 48C12, which belongs to its hydrophobic extracellular part. There are no analogues in microbial rhodopsins for Glu202, which thus may be a key determinant of the subfamily 2 protein function. A highly conserved Pro172, which makes a π-bulge in helix E of 48C12, also characteristic only for HeRs, is absent in group 2, however, proline is present in position 168 (helix E) of 48C12 in almost all its members. This alteration may change the shape of helix E and affect the folding of the protein. The retinal binding pocket in HeRs of group 2 is extremely different from that of other subfamilies, especially due to the presence of positively charged His residues in positions 162 and 166 of the reference protein 48C12. Analogues of Asn138 are also absent in group 2.

*Subfamilies 3, 4, 5,* have variations from 48C12 in the retinal binding pocket. Particularly, methionine and asparagine in subfamily 3 are placed in the positions of Gln213 and Ile142 of 48C12, respectively. The same asparagine is present in groups 4 and 5, however, it alternates with asparagine in the position of Asn138, thus only the Asn residue is present near the β-ionone ring of the retinal.

*Subfamilies 7, 8 and 9* have a very interesting feature of conservative Tyr in position 202 of 48C12. Asn is present in the position of Ile142 of 48C12 in all members of groups 8 and 9 and in some representatives of subfamily 7. Group 9 also has no analogues of Asn138 of 48C12.

*“Unsorted proteins” group* includes the most different heliorhodopsins. These proteins presumably maintain the polar cavity in the cytoplasmic part, however, its surroundings are varied. Most interesting, in subgroup U1, histidine is present in the position of Asn16 in addition to two histidines in the positions of His23 and His80 of 48C12. Moreover, at the extracellular side two glutamates are present in all members of subfamily U1 in the positions of Leu73 and Ile116 of 48C12. Glutamate is also found in the positions of Pro172 (subfamily U2), Val69 and Leu202 (subfamily U8) of 48C12. Positively charged residues also appear in the extracellular side of the members of subfamilies U1, U6, U7, and U11, such as His residues in the position of Leu73 and Arg and Lys residues in the position of Leu253 of 48C12. The positions of 48C12, whose analogues in other HeRs are occupied by unusual charged or polar residues and possible variants of those residues are shown in Fig. 6. These charged amino acids, especially located in the extracellular part of the proteins, may be crucial for the functions of those HeRs.

Subfamilies 3, 5, 6, 8, U1, U2, U3, U4, U5, U6 and U12 consist exclusively of bacterial proteins. Subfamilies 4, 7, 9, and U13 represent archaeal HeRs (except one bacterial protein from subfamily 4 and one from subgroup U13), mostly *Euryarchaeota*, but subfamily 7 also has members of *Asgard* and *TACK* groups. Subfamilies U8, U9, U11 comprise proteins of eukaryotic origin.

## Discussion

### Molecular mechanisms and biological function(s) of HeRs

A surprise of the first works on HeRs (the studies of the 48C12) is that the attempts to identify amino acids playing the roles of primary proton acceptor and proton donor to the RSB failed^20, 21^. Such amino acids are key functional determinants in all known rhodopsins. Another important fact is that Pushkarev *et al*. did not observe any translocation of the proton (an ion) through the protein to its polar surfaces. High resolution crystallographic structures of 48C12 HeR which represents the most abundant subfamily of HeRs (195 out of 479 currently known unique sequences), were solved at 1.5 Å resolution with the crystals obtained at pH 8.8 and 4.3, respectively. The structures correspond to two different forms of the protein. Both structures show a remarkable difference between HeRs and all the known rhodopsins. The retinal binding pocket, those parts of the cytoplasmic and extracellular regions of the protein which are determinants of the function of the known rhodopsins are also different. There is no analogue to this protein among type 1 (microbial) and type 2 (visual) rhodopsins.

In the cytoplasmic part of the protein at pH 8.8 there is a large cavity (‘active site’) placed close to the RSB and surrounded by highly conservative charged amino acids His23, His80, Arg104, Glu107, Glu230, the protonated RSB, by polar residues Asn16, Tyr92, Asn101, Tyr108, Ser237 and filled with 6 water molecules. The amino acids and the RSB are interconnected by an extensive hydrogen network mediated by the water molecules (Fig. 2). Notably, there are two pathways from the cavity to the bulk. From one of the sides the cavity is separated from the bulk by only the Asn101 residue at the surface of the protein at the level of the hydrophobic/hydrophilic interface (Fig. 4, Fig.S10). From the other side – by Arg104, found in almost all rhodopsins as a major gate between the RSB and the bulk.

A major difference between the two structures is that at lower pH the ‘active site’ comprises planar triangle shaped molecule in the cavity. Remarkably, several mentioned above residues (namely His23, His80, Arg104, Glu107, Tyr108, Ser237) were subjected to alanine substitution^21^ which in all cases led to the changes of absorption spectra. This result supports the presence of a strong interaction of the active site with the RSB. In its turn, it means that isomerization of the retinal modifies the properties of the ‘active site’ since the base is directly connected to the cavity through Glu107 via a hydrogen bond (Fig. 3B, Fig. 5D).

The structure suggests also why previous attempts to identify a proton acceptor and a proton donor failed^20, 21^. One of the reasons is that one of the possible amino acid candidate for these roles, for instance, Glu230 which is a key member of the active site, was overlooked in previous studies because of poor prediction of the protein topology in the membrane (Fig. 1 in ref ^21^). However, the structure and additional mutational analysis suggest that the cluster of water molecules in the ‘active site’ plays a role of a reservoir for the dissociated from the RSB proton. Importantly, the RSB proton transfer to the hydrophobic extracellular part of the protein upon isomerization of the retinal seems to be problematic due to high free energy penalty. We suppose that the RSB proton dissociates upon isomerization of the retinal but does not leave the inner cavity during photocycle. This proton might interact with the substrate in the reaction H^+^ + substrate^-^ _▭_ reduced substrate like in carbon fixation which is known as one of the most important biosynthetic processes in biology^33^.

### Biological role of HeR Subfamily 1

At this point it is difficult to establish the primary role of HeR even of the best known Subfamily 1. Pushkarev *et al.* suggested that HeRs may function as sensory proteins^20^. This conclusion was solely based on two observations. First, the authors did not detect any ion translocation activity of the protein (under experimental conditions corresponding to pH 8.1). Second, the photocycle of the protein (measured at pH 8.5) was several seconds long which is a characteristic of sensory rhodopsins^3^. Along these lines, we noted above that heliorhodopsin possesses a hydrogen bond-forming aromatic amino acid Trp246 that faces the membrane and that might change its conformation under illumination, similarly to Tyr199 in *Np*SRII^14, 31^. Usually the genes of sensory rhodopsins have a transducer gene transducer located nearby and often co-transcribed ^34, 35^. At present, two distinct types of sensory rhodopsins are known: SRII-like photoreceptors utilize transmembrane chemoreceptor-like transducer proteins, whereas *Anabaena* sensory rhodopsin (ASR) utilizes a soluble transducer protein that dissociates from ASR upon illumination^3^. In case of subfamily 1, however, no conserved proteins, which could potentially be signal transducers, could be detected in the genomic neighborhood. On the other hand, in some microbes (aquatic actinobacteria such as the marine *Ca.* Actinomarina) that most often contain heliorhodopsins, their genes are surrounded by two large clusters of Nuo genes (Fig. S18), which products are key proteins in respiratory chains. At this point it is unclear what it might mean. However, the ability of HeRs to bind the triangle anions like carbonate in the ‘active site’ suggests the possibility of its involvement in carbon fixation.

The analysis of the presence/absence of HeRs in monoderm and diderm representatives of the Tara Oceans and 25 freshwater lakes metagenomes led to the conclusion that heliorhodopsins were absent in diderms, confirming their absence in cultured *Proteobacteria*. Judging by a specific semipermeability of outer membranes of diderms they proposed a role of HeRs in light-driven transport of amphiphilic molecules^22^. However, the structures of 48С12 do not support such function for HeRs. Moreover, according to the literature data we conclude that, in fact, there is no clear evidence of the HeRs presence only in monoderms^36^. For example, some of the proteins of subfamily 1 are originating from *Bacteroidetes*, *Gemmatimonadetes* and from *Proteobacteria*, all of which are assumed to be diderms. Some HeRs from other subfamilies are found in the phyla *Thermotogae* and *Dictioglomy* which have also diderm cells.

Although at this point we cannot provide a definitive role for HeR we would like to advance a hypothesis. In many (if not most cases) rhodopsins are found in pelagic microbes living in the photic zone of aquatic habitats (freshwater or marine). They appear in microbes that often contain also a classical rhodopsin, typically a proton pump, providing the cell with unlimited energy as long as there is light. The transfer of the retinal proton to the interior of the cell, likely reducing a molecule of carbonate or nitrate might act like cyanobacterial (or plant) photosystem II transforming light energy into reducing power to form precursors of cell biomass. This would transform the microbes containing the two kind of rhodopsins in primary producers like cyanobacteria, and would help explaining the extraordinary success of some of them, such as the actinobacteria that are the most abundant microbes in most photic freshwater habitats. Further structure-guided functional studies are necessary to clarify the biological role (roles) of this completely different family of rhodopsins.

## Materials and Methods

### Protein expression and purification

The gene of helirhodopsin 48C12 (UniProt ID A0A2R4S913; NIH Genbank AVZ43932.1) was synthesized *de novo* and optimized for expression in *E.coli* with Thermo Fisher Scientific GeneOptimizer service. The optimized gene was introduced into Staby™Codon T7 expression plasmid system (Delphi Genetics, Belgium) via NdeI and XhoI (Thermo Fisher Scientific) that led to the addition of 6×His tag to the C-terminus of the gene. The resulting plasmid DNA was sequenced (Eurofins Genomics, Germany) and used to transform *E.coli* C41 strain.

The protein expression procedure is adopted from ref ^37^ but slightly modified. The culture was cultivated at 37 °C in the auto induction media (1% w/v Trypton, 0.5% w/v Yeast extract, 0.5% w/v glycerol, 0.05% w/v glucose, 0.2% w/v lactose, 10 mM (NH_4_)_2_SO_4_, 20 mM KH_2_PO_4_, 20 mM Na_2_HPO_4_, adjusted pH 7.8) containing 150 µg/ml of ampicillin antibiotic to OD_600_=0.8. After, the cultivation temperature was decreased to 26 °C with subsequent addition of 150 µg/ml ampicillin, 20 µM all-trans Retinal (solubilized in Triton X-100 detergent) and 0.1mM IPTG and the culture was grown overnight. The concentration of antibiotic after induction was maintained with addition of extra 150 µg/ml each two hours.

The cells were then collected and disrupted at 20000psi with M-110P homogenizer (Microfluidics) in the buffer containing 30mM Tris-HCl pH 8.0, 0.3M NaCl, 0.04% Triton X-100, 50 mg/L DNase I (Sigma-Aldrich) and cOmplete® protease inhibitor cocktail (Roche). The total cells’ lysate was ultracentrifugated at 120000 rcf. Then, membranes were isolated and dispensed in the same buffer without DNAse (whit addition of 1% w/w DDM detergent and 5mM all-trans retinal) and stirred overnight at 4 °C.

The non-soluble fraction was separated by ultracentrifugation at 120,000 rcf for 1 h at 4 °C. The resulting soluble protein mixture was loaded to Ni-NTA resin (Cube Biotech). The column with loaded resin was washed with 3 CV of washing buffer WB1 (30 mM Tris-HCl pH 8.0, 0.3 M NaCl, 10 mM Imidazole, 0.05% Triton, 0.2% DDM) and washing buffer WB2 (30 mM Tris-HCl pH 8.0, 0.3 M NaCl, 50 mM Imidazole, 0.05% Triton, 0.2% DDM). Then, heliorhodopsin was eluted with EB buffer (30 mM Tris-HCl pH 7.8, 0.3 M NaCl, 250 mM L-Histidine (AppliChem), 0.05% Triton, 0.1% DDM). The eluted protein mixture was subjected to the size-exclusion chromatography column Superdex200 Increased 10/300 GL (GE Health Care Life Sciences) pre-equilibrated with SEC buffer (30 mM Tris-HCl, 50 mM NaPi pH 7.8, 300 mM NaCl, 0.5 mM EDTA, 2 mM 6AHA (6-Aminohexanoic Acid), 0.075% DDM). The fractions were analyzed and those containing the 48C12 rhodopsin with peak ratio of ∼1.25 and lower were collected and protein was concentrated to 20 mg/ml with 50 kDa concentration tubes at 5000 rcf and flash-cooled with liquid nitrogen.

### Flash photolysis setup

The laser flash photolysis was similar to that described by Chizhov et al. ^38–40^ with minor differences. The excitation system consisted of Nd:YAG laser Q-smart 450mJ with OPO Rainbow 420-680nm range (Quantel, France). Samples were placed into 5×5mm quartz cuvette (Starna Scientific, China) and thermostabilized via sample holder qpod2e (Quantum Northwest, USA) and Huber Ministat 125 (Huber Kältemaschinenbau AG, Germany). The detection system beam emitted by 150W Xenon lamp (Hamamatsu, Japan) housed in LSH102 universal housing (LOT Quantum Design, Germany) passed through pair of Czerny–Turner monochromators MSH150 (LOT Quantum Design). The received monochromatic light was detected with PMT R12829 (Hamamatsu). The data recording subsystem represented by a pair of DSOX4022A oscilloscopes (Keysight, USA). The signal offset signal was measured by one of oscilloscopes and the PMT voltage adjusted by Agilent U2351A DAQ (Keysight). The absorption spectra of the samples were measured before and after each experiment on Avaspec ULS2048CL fiber spectrophotometer paired with AVALIGHT D(H)S Balanced light source (Avantes, Netherlands).

### Preparation of samples for flash photolysis

The wild type protein sample (WT) for flash photolysis assay was purified in the same manner as for crystallization but with increased from 0.3 M to 0.6 M NaCl concentration on each purification step. The purified WT protein was 100x diluted in buffer containing 30 mM HEPES pH 7.0, 1 M NaCl, 2% DDM to concentration of ca. 0.5 mg/ml. The measurement was performed in the following way. The 350 ul sample was placed into the 5 mm light path cuvette and the temperature of the sample was set to 20⁰C. After, the protein sample was exposed to 6 ns pulse of 3.5 mJ mean (standard deviation 6% on 1000 pulses) @545 nm. The transient absorption changes data was recorded (in 350-700nm light range; step 10nm) from 1mks up to 5 sec with two oscilloscopes with overlapping ranges (range ratio 1:1000) and averaged for 20 pulses for each wavelength. The data compression reduced the initial number of data points per trace to ca. 900 points. The samples of E230Q and E149Q mutant proteins were prepared without purification similar to ^21^ with modification. The E.coli C41 cells were disrupted at 20000psi with M-110P homogenizer in buffer containing 30mM Tris-HCl pH 8.0, 1 M NaCl and DNase I and non-soluble fraction was sedimented at 120000 rcf. The 5 g of the membranes were then washed and resuspended in 20ml of the buffer containing 30mM HEPES pH 7.0, 1M NaCl. After homogenization, the 2% DDM were added to 1.6ml of suspension and the sample was incubated for 30 min at 4 ⁰C. Later, samples were applied to the centrifugation (for 10 min at 4 ⁰C, 15000 rcf) and supernatant was collected for characterization. The flash photolysis measurement of E230Q/E149Q mutant-containing samples was performed at 400-610nm (step 70nm; each reading averaged for 20 pulses) at 20 ⁰C using 6 ns excitation pulses of 3.5 mJ @545 nm.

### Crystallization

The crystals were grown with an *in meso* approach^14, 41^, similar to that used in our previous work^16, 23^. The solubilized protein in the crystallization buffer was mixed with pre-melted at 42°C monoolein (Nu-Chek Prep) to form a lipidic mesophase. The 150 nl aliquots of a protein–mesophase mixture were spotted on a 96-well LCP glass sandwich plate (Marienfeld) and overlaid with 500 nL of precipitant solution by means of the NT8 crystallization robot (Formulatrix). The best crystals of the violet form were obtained with a protein concentration of 20 mg/ml and the precipitant solution of 2.0 M Ammonium Sulfate, 0.1 M Tris-HCl pH 8.8. For the blue form the best crystals were grown with the same protein concentration of 20 mg/ml and the precipitant solution of 2.0 M Ammonium Sulfate, 0.1 M Sodium acetate pH 4.3. The crystals were grown at 20°C to observable size in two weeks for the both types. The rhombic-shaped crystals reached 150 μm in length and width with maximum thickness of 20 μm. Crystals of the both forms were incubated for 5 min in cryoprotectant solution (2.0 M Ammonium Sulfate, 0.1 M Tris-HCl pH 8.8 for the violet form and 2.0 M Ammonium Sulfate, 0.1 M Sodium acetate pH 4.3 for the vlue form supplied with 20% (w/v) glycerol) before harvesting. All crystals were harvested using micromounts (MiTeGen), and were flash-cooled and stored in liquid nitrogen. Absorption spectra from the 48C12 crystals were measured at ID29s beamline of European Synchrotron Radiation Facility (ESRF), Grenoble, France at 300 K^42^.

### Collection and treatment of diffraction data

X-ray diffraction data were collected at Proxima-1 beamline of the SOLEIL, Saint-Aubin, France at 100 K, with an EIGER 16M detector and at P14 beamline of the PETRAIII, Hamburg, Germany France at 100 K, with an EIGER 16M detector. We processed diffraction images with XDS^43^ and scaled the reflection intensities with AIMLESS from the CCP4 suite^44^. The crystallographic data statistics are presented in Extended Data Table 1. The molecular replacement search model was generated by RaptorX web server^45^ based on the ESR structure (PDB ID 4HYJ^6^). Initial phases were successfully obtained in P2_1_ space group by the molecular replacement using phenix.mr_rosetta^46^ of the PHENIX^47^ suite. The initial model was iteratively refined using REFMAC5^48^, PHENIX and Coot^49^. The cavities were calculated using HOLLOW^25^. Hydrophobic-hydrophilic boundaries of the membrane were calculated using PPM server^50^.

### Bioinformatics analysis

Multiple amino acid alignment was performed using Clustal Omega algorithm^51^. Heliorhodopsins database was downloaded from InterPro^32^ and merged with database provided in original article^20^. Phylogenetic tree was constructed and classes identified using iTOL server software version 4.3.2 ^52^. For removing those proteins above certain similarity threshold we used CD-HIT suite^53^. Cut-off similarity threshold is always specified. Calculations of conservative amino acids were performed using an in-house C# application written using Visual Studio Community 2017. Most conservative regions were identified and normalized results were visualized using in-house Wolfram Mathematica Notebooks. Genome sequence of the single-amplified genome AG-333-G23, belonging to the marine Ca. Actinomarinales group, was downloaded from the NCBI database (Biosample: SAMN08886063). Encoded genes were predicted using Prodigal v2.6^54^. tRNA and rRNA genes were predicted using tRNAscan-SE v1.4^55^, ssu-align v0.1.1^56^ and meta-rna^57^. Predicted protein sequences were compared against the NCBI nr database using DIAMOND^58^, and against COG^59^ and TIGFRAM^60^ using HMMscan v3.1b2^61^ for taxonomic and functional annotation. A custom database containing both type I and type III rhodopsins^20^ was used to identify putative homologs. Resulted significative genes (HMMscan, E-value 1e-15) were then confirmed by determining the secondary structure and the presence of domains with InterPro^32^.

## General

We thank O. Volkov and A.Yuzhakova for technical assistance. We acknowledge the Structural Biology Groups of the Swiss Light Source (SLS, Villigen, Switzerland), SOLEIL (Saint-Aubin, France) and PETRAIII (Hamburg, Germany) for granting access to the synchrotron beamlines. We are grateful to A. Royant (ID29S Cryobench laboratory, ESRF) for help with collection of the 48C12 crystals optical absorption spectra.

## Funding

This work was supported by the common program of Agence Nationale de la Recherche (ANR), France and Deutsche Forschungsgemeinschaft, Germany (ANR-15-CE11-0029-02/FA 301/11-1 and MA 7525/1-1) and by funding from Frankfurt: Cluster of Excellence Frankfurt Macromolecular Complexes by the Max Planck Society (to E.B.) and by the Commissariat à l’Energie Atomique et aux Energies Alternatives (Institut de Biologie Structurale)–Helmholtz-Gemeinschaft Deutscher Forschungszentren (Forschungszentrum Jülich) Special Terms and Conditions 5.1 specific agreement. This work used the platforms of the Grenoble Instruct-ERIC center (ISBG; UMS 3518 CNRS-CEA-UJF-EMBL) within the Grenoble Partnership for Structural Biology (PSB). Platform access was supported by FRISBI (ANR-10-INBS-05-02) and GRAL, a project of the University Grenoble Alpes graduate school (Ecoles Universitaires de Recherche) CBH-EUR-GS (ANR-17-EURE-0003). The reported study was funded by RFBR and CNRS according to the research project № 19-52-15017. F.R.-V. thanks the grant ‘VIREVO’ CGL2016-76273-P [AEI/FEDER, EU], (co-funded with FEDER funds).

## Author contributions

DV made cloning, expression, purification and functional characterization of the protein and cloning, expression and flash-photolysis investigation of E230Q and E149Q mutants; RA performed its crystallization; AA made bioinformatics analysis; KK did spectroscopic studies of the crystals; KK and RA collected the diffraction data with the help of GB; KK solved the structures; IG helped with structure solving, refinement and analysis; AA performed sequence alignment analysis; J.M.H.-M. performed genome analysis of Ca. Actinomarinales; GB, AP, AR, VB, EB and F.R-V. contributed to data analysis and editing the manuscript; VG designed and supervised the project; KK and VG analyzed the results and prepared the manuscript with input from all the other authors.

## Competing interests

Authors declare no competing interests

## Supplementary Materials

**Fig. S1.**
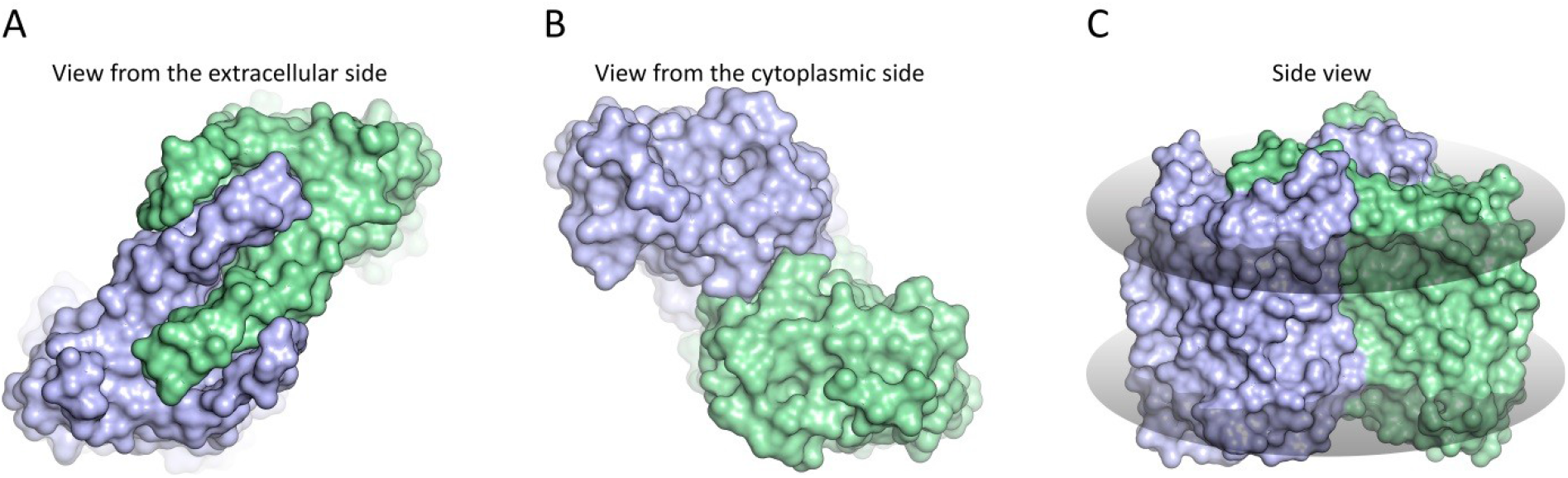
Surface representation of 48C12 dimer. **A.** View from the extracellular side. **B.** View from the cytoplasmic side. **C.** Side view in the membrane.

**Fig. S2.**
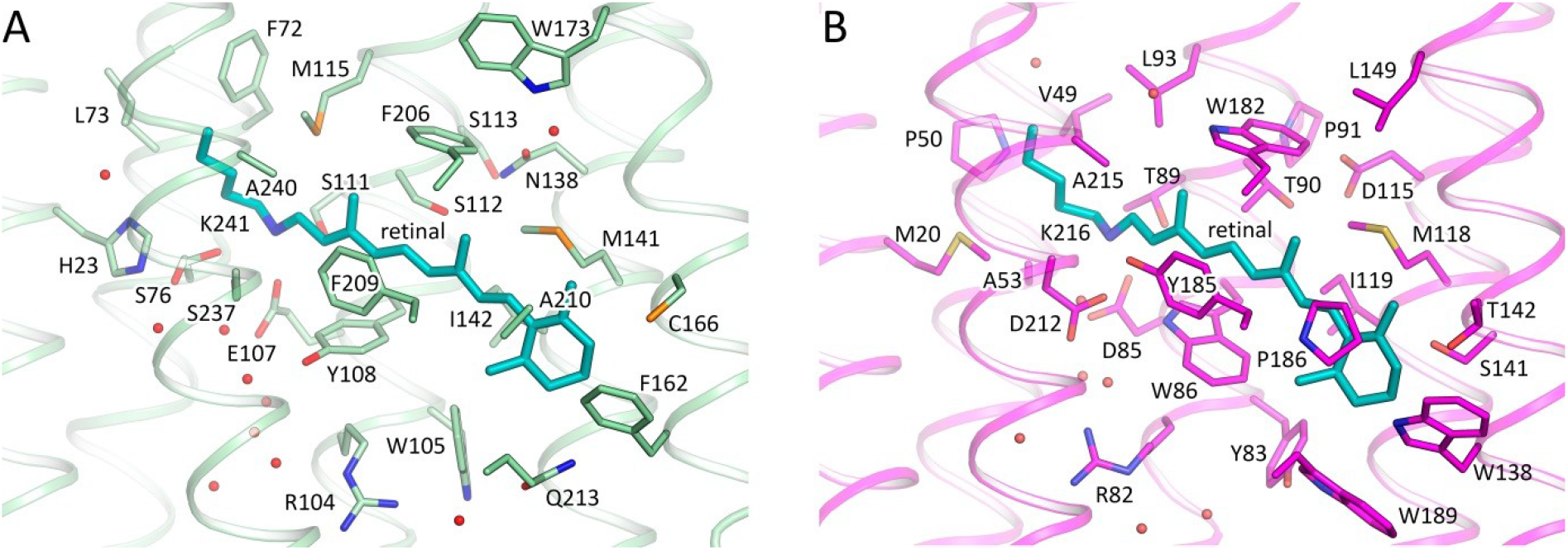
Retinal binding pocket of. **A.** 48C12**. B.** bR. Cofactor retinal is colored teal.

**Fig. S3.**
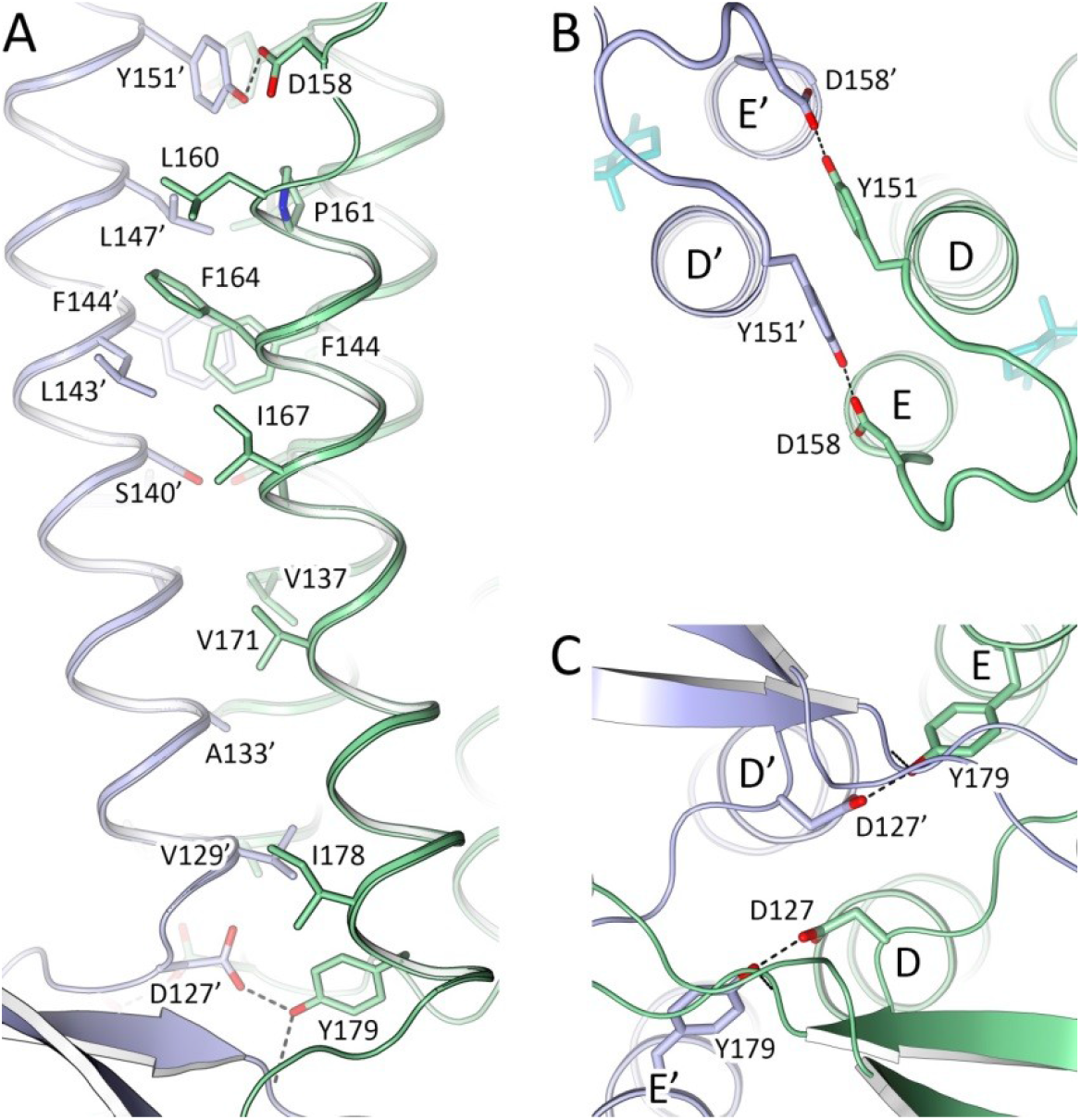
Dimerization interface of 48C12. **A.** Side view of the oligomerization interface. Extracellular side is on the bottom. **B.** View from the cytoplasmic side. **C.** View from the extracellular side. The strongest contacts by hydrogen bonding are shown with dashed lines.

**Fig. S4.**
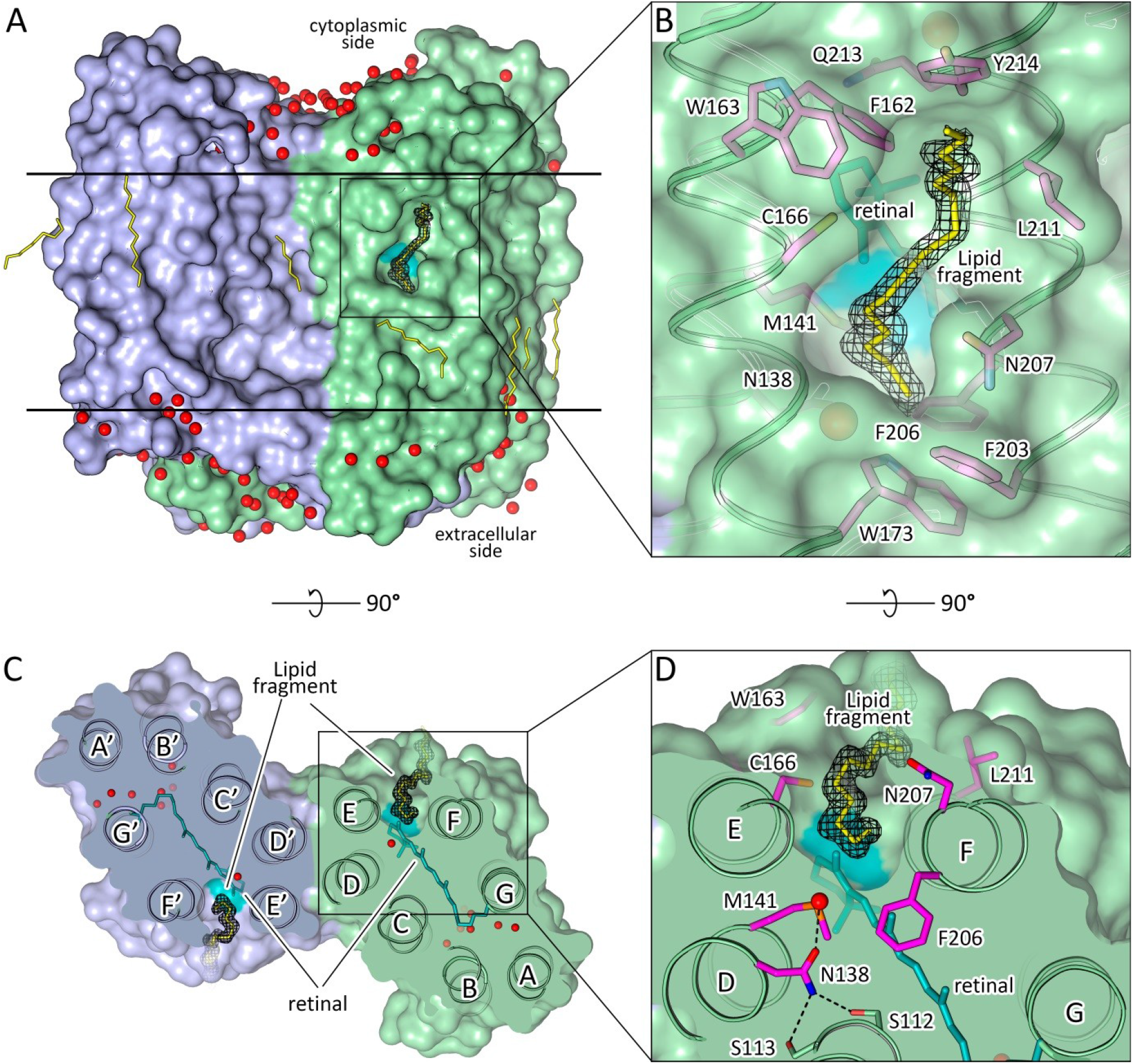
Lipid molecule permeating into 48C12 protomer. **A.** Surface view of the 48C12 dimer in the membrane with lipid molecules. Cofactor retinal is colored teal. Membrane core boundaries are shown with black lines. **B.** Detailed side view of the lipid molecule inside the protomer. Residues, comprising the pocket for the molecule are colored violet. **C.** Section view from the extracellular side. Lipid molecule is deepened into the protomer between helices E and F. **D.** Detailed section view from the extracellular side. Hydrogen bonds are shown with black dashed lines. 2FoFc electron density maps around lipid molecule are contoured at the level of 1.5σ

**Fig. S5.**
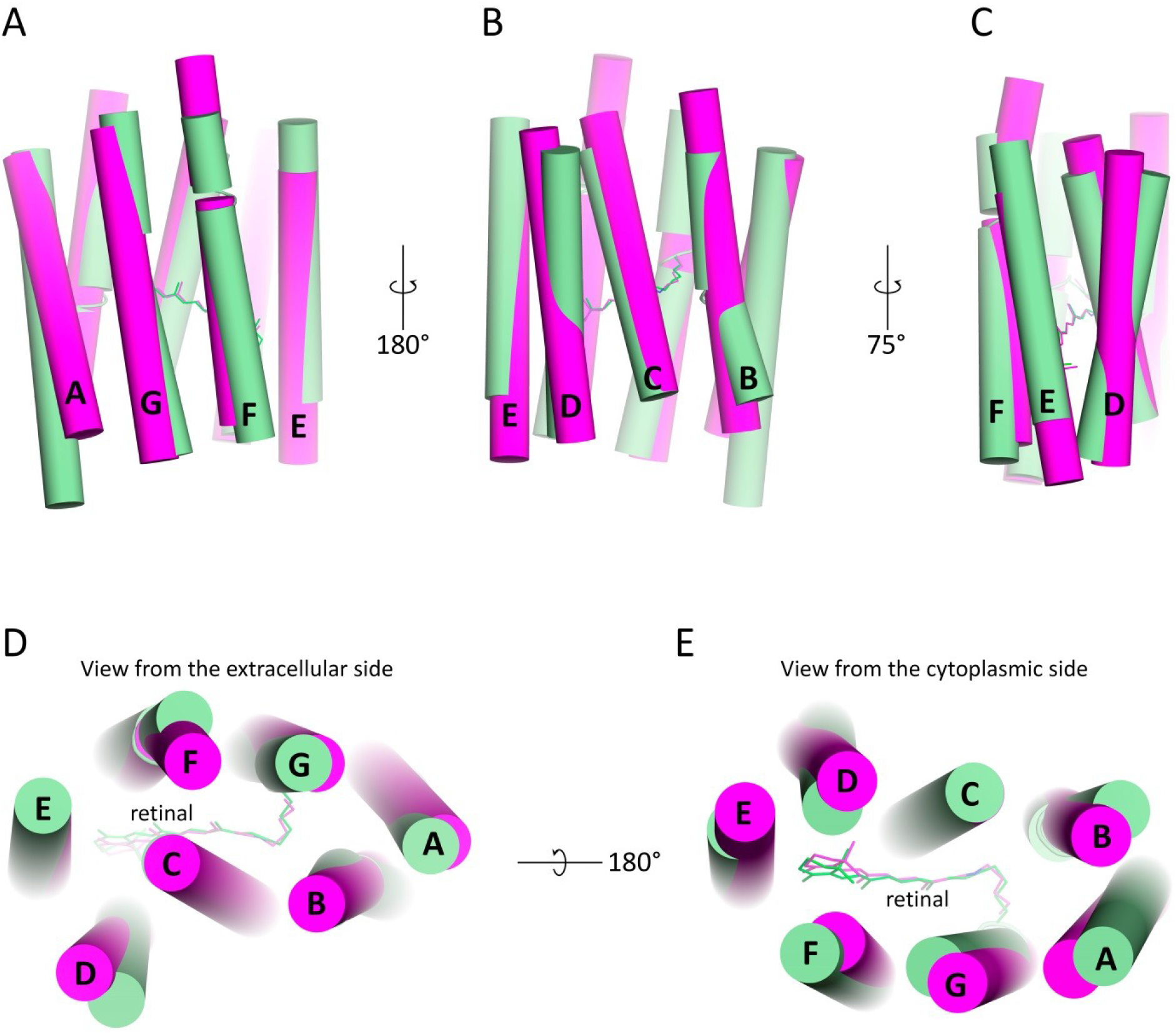
Structure alignment of 48C12 (green) and bR (purple, PDB ID 1C3W). **A.** Side view from the side of helices F and G. **B.** Side view from the side of helices B and C. **C.** Side view from the side of helices D and E. **D.** View from the extracellular side. **E.** View from the cytoplasmic side.

**Fig. S6.**
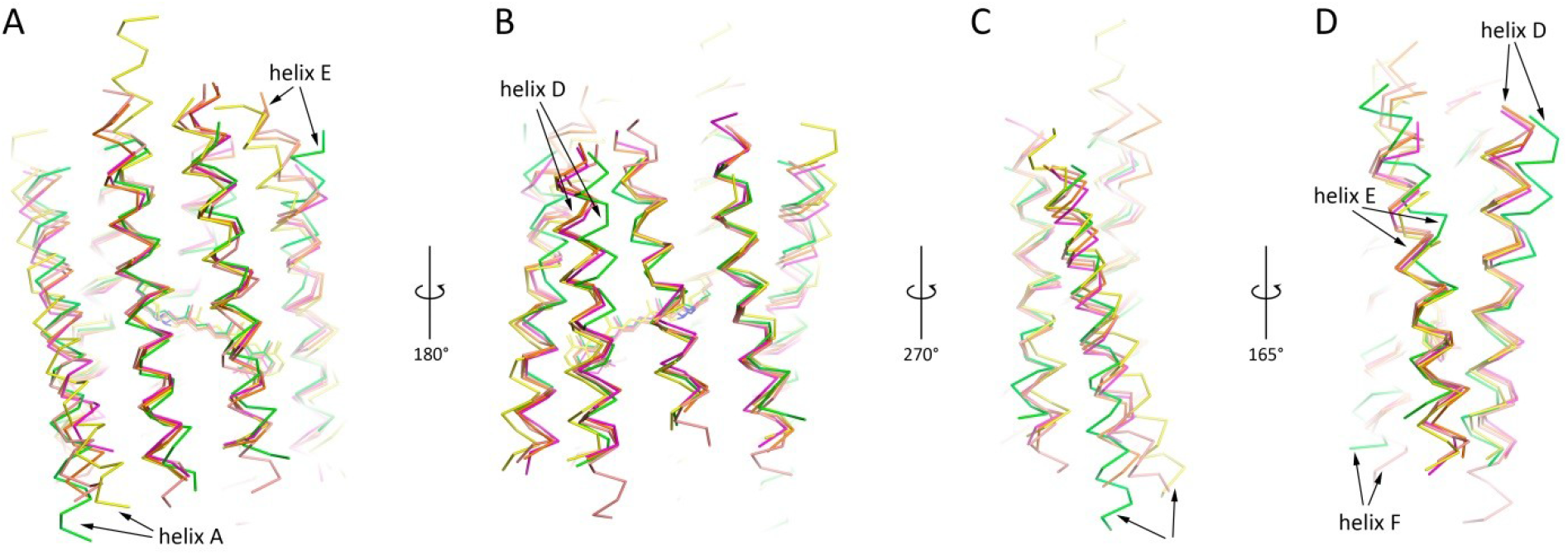
Structure alignment of 48C12 (green), bR (purple, PDB ID 1C3W), NpSRII (orange, PDB ID 3QAP), ChR2 (yellow, PDB ID 6EID) and ESR (salmon, PDB ID 4HYJ). **A.** Side view from the side of helices F and G. **B.** Side view from the side of helices B and C. **C.** Side view from the side of helix A. **D.** Side view from the side of helices D and F. Black arrows indicate the most significant differences in helices location between 48C12 heliorhodopsin and representatives of type 1 rhodopsins. Extracellular side for 48C12 is at the top, cytoplasmic side is at the bottom of each figure section.

**Fig. S7.**
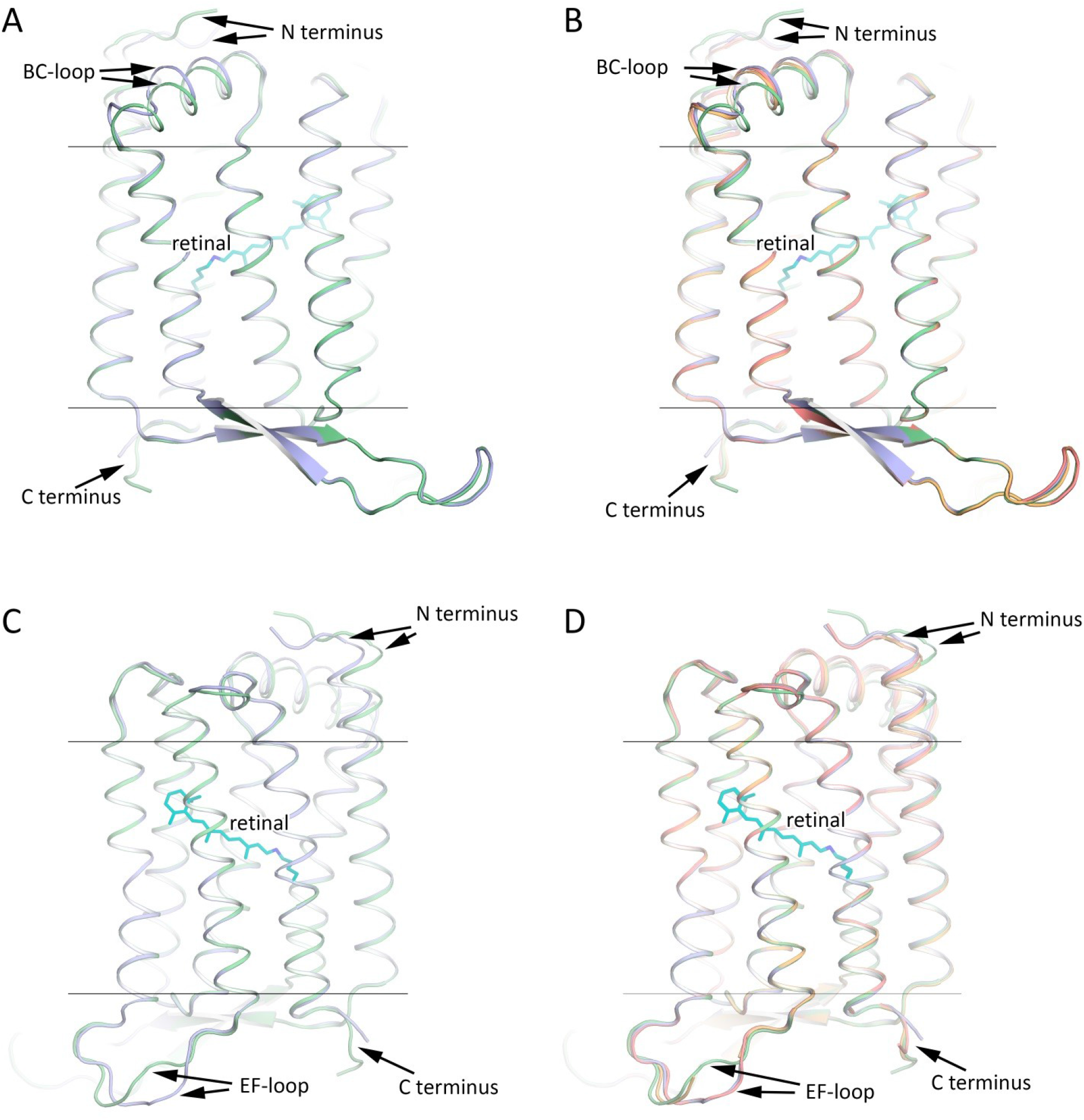
Structure alignment of the backbones of 48C12 protomers. Alignment of protomer A (green) and protomer B (blue) at neutral pH. Side view from the side of helices B and C (**A**) and from the side of helices F and G (**C**). Alignment of protomer A (green) and protomer B (blue) at neutral pH and protomer A (orange) and protomer B (red) at acidic pH. Side view from the side of helices B and C (**B**) and from the side of helices F and G (**D**). Extracellular side for 48C12 is at the top, cytoplasmic side is at the bottom of each figure section.

**Fig. S8.**
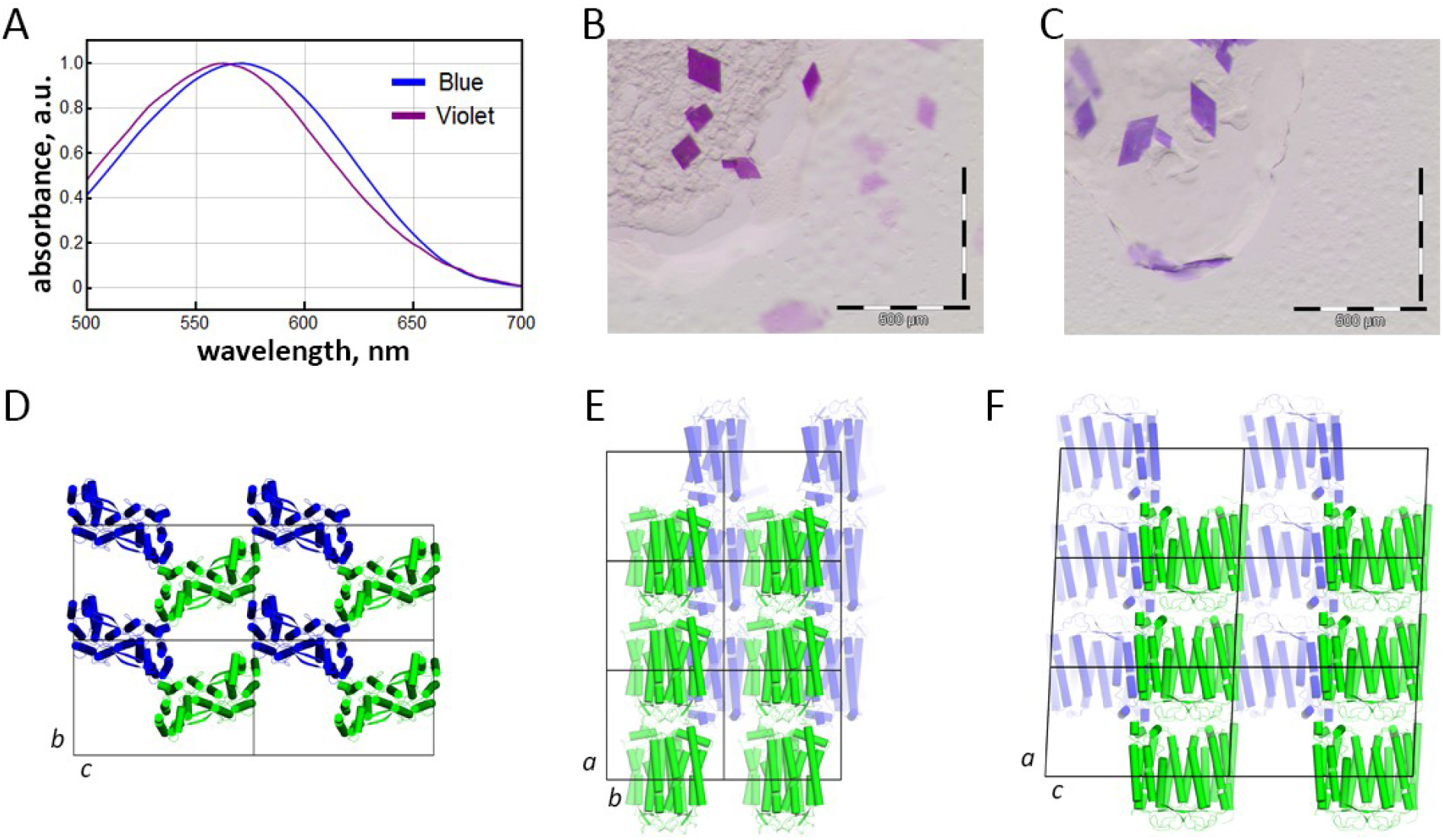
Crystals and crystal packing of 48C12. **A.** Absorption spectra from 48C12 crystals of violet and blue forms. **B.** Example of violet crystals of 48C12. **C.** Example of violet crystals of 48C12. **D-F.** Crystal packing of both violet and blue forms of 48C12.

**Fig. S9.**
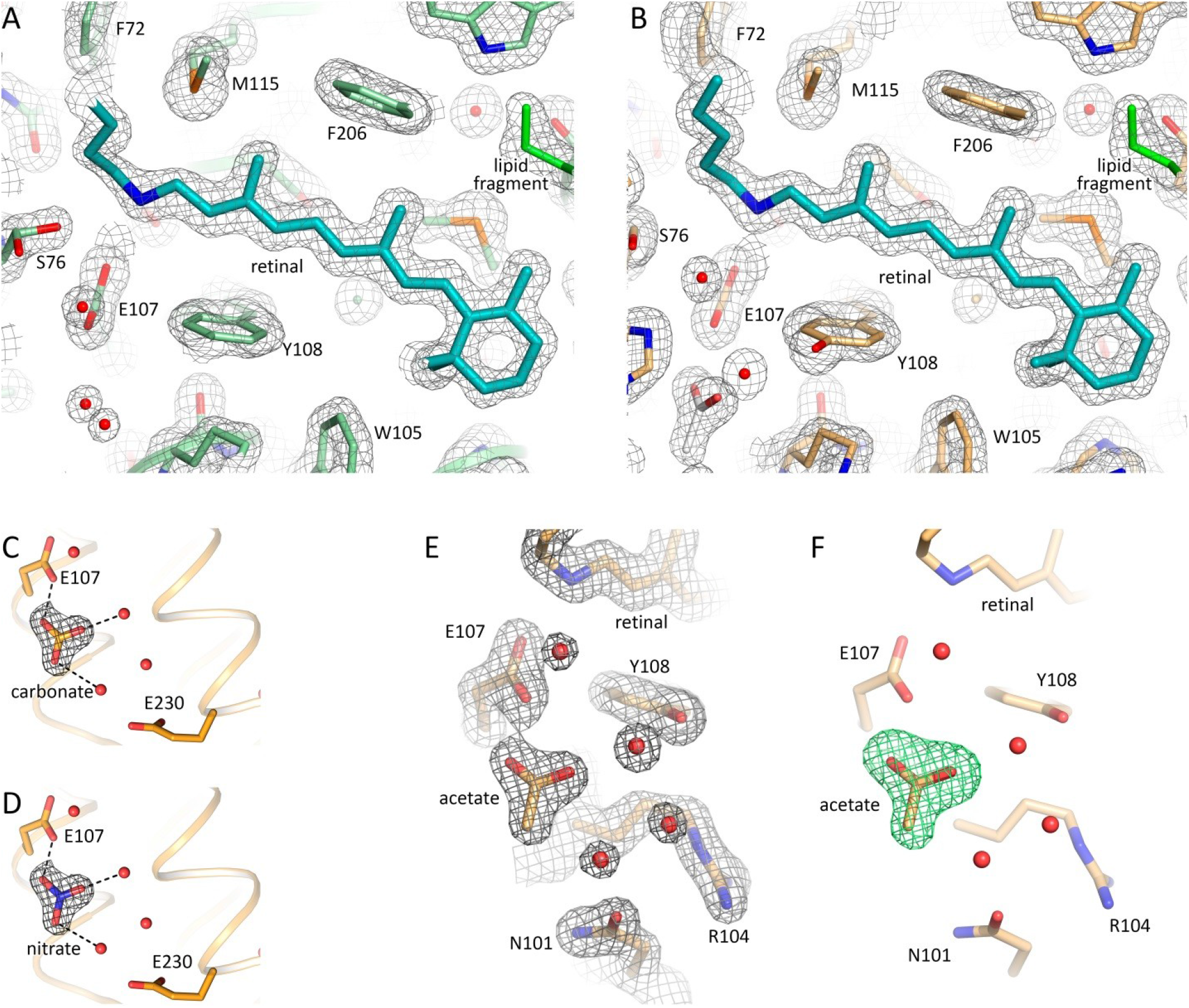
Examples of electron density maps of 48C12 model. A, B. 2Fo-Fc electron density maps near the retinal of the chain A of violet and blue forms, respectively. The maps are contoured at the level of 1.5σ. **C, D.** Examples of 2Fo-Fc electron density maps of the active site of the protein after hypothetical fitting of the triangle difference electron densities with carbonate and nitrate molecules, respectively. Carbonate and nitrate molecules were placed for modelling at the position of the acetate in the blue form of 48C12. Putative hydrogen bonds are shown with black dashed lines. The maps are contoured at the level of 1.5σ. **E.** 2Fo-Fc electron density maps of the active site and acetate molecule in the blue form of the 48C12 heliorhodopsin. The maps are contoured at the level of 1.5σ. **F.** Simulated annealing omit Fo-Fc difference electron density maps calculated using *phenix.polder* omitting the acetate molecule. The maps are contoured at the level of 3σ.

**Fig. S10.**
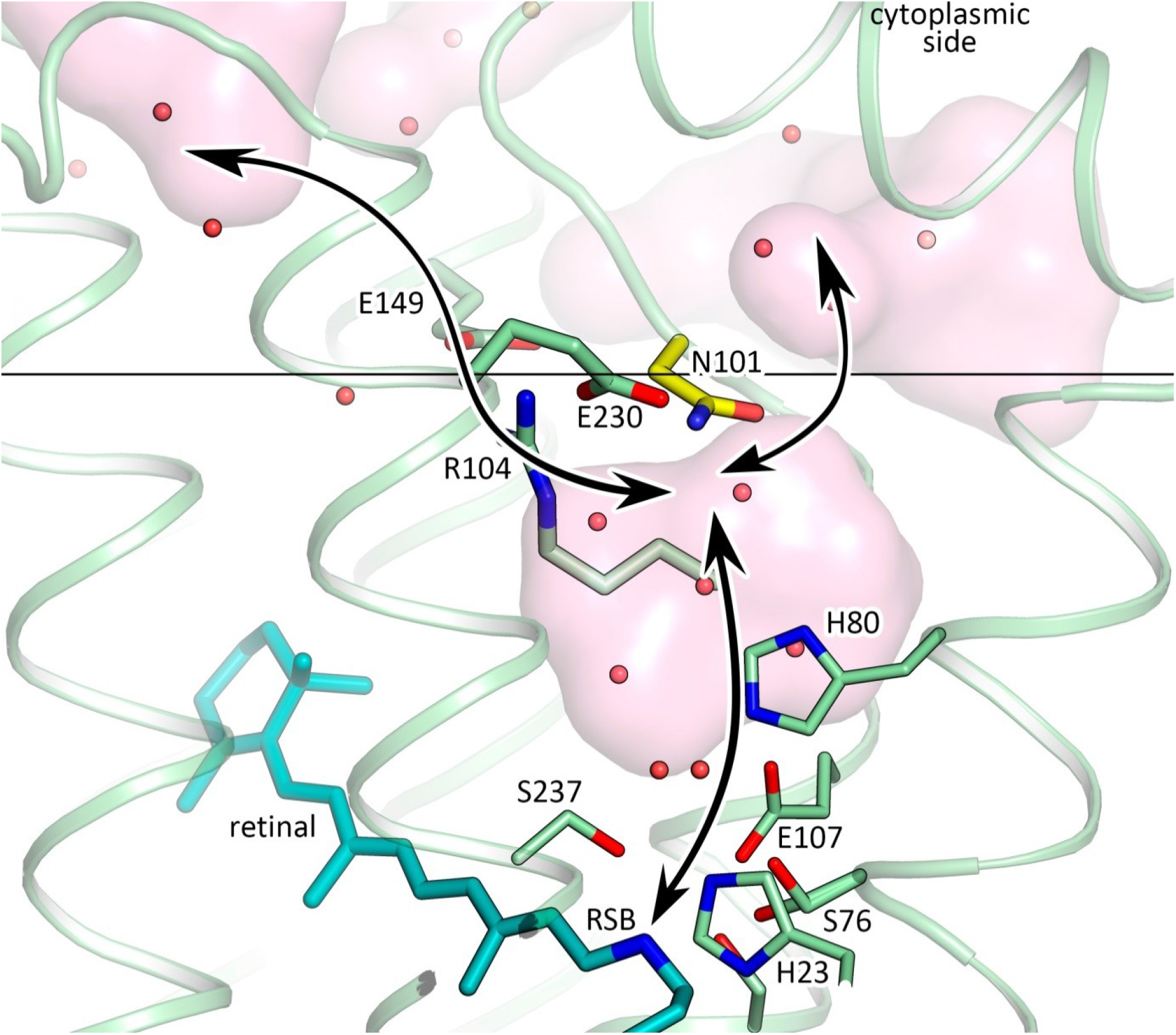
Putative pathway of the proton in 48C12 heliorhodopsin. Black arrows indicate the putative way of the protons. Membrane core boundary at the cytoplasmic side is shown with black line. Asn101 residue is colored yellow.

**Fig. S11.**
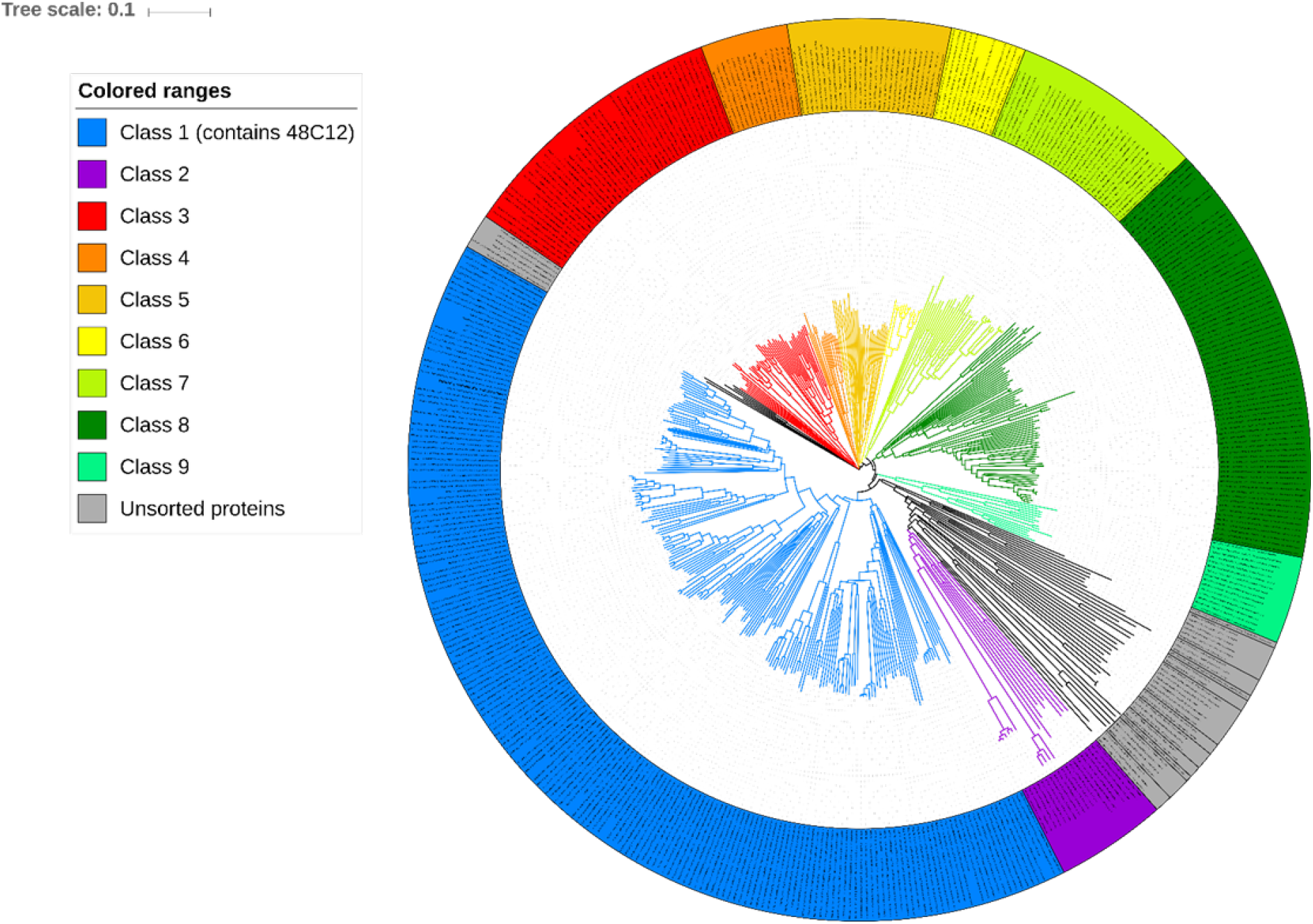
Phylogenetic tree for heliorhodopsins. Clades whose average branch length distance to leaves if below 0.397 are collapsed into classes. Classes containing more than 10 proteins assigned with index number and color (see legend). Classes containing less than 10 proteins assigned with gray color and “Unsorted proteins” label. Analyzed base of heliorhodopsins consists of proteins from original article^20^ and proteins predicted with InterPro.

**Fig. S12.**
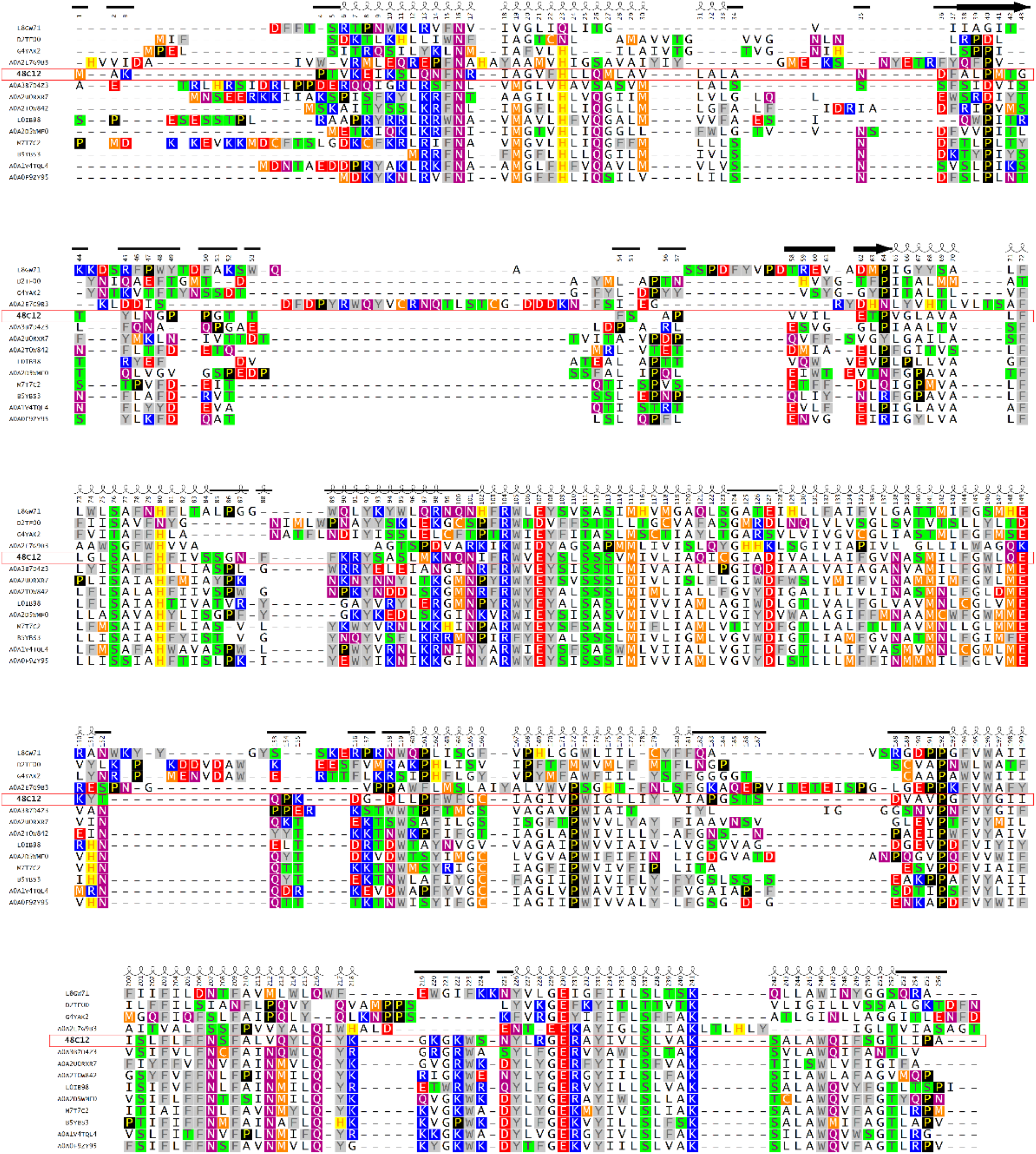
Sequence alignment of heliorhodopsins of different classes (Heliorhodopsin 48C12 numbering). Proteins represent a different part of a phylogenetic tree and different domains of life. Heliorhodopsin 48C12 (A0A2R4S913 Uniprot ID), A0A3B7D4Z3 (Uniprot ID) and A0A2D5WMF0 - Bacteria>Terrabacteria group>Actinobacteria both, L8GW71 –Eukaryota>Amoebozoa, A0A2E7G9B3 - Bacteria>Proteobacteria, D2TF00 and G4YAK2 – Viruses both, L0IB98 - Archaea>Euryarchaeota, A0A2T0W842 - Bacteria>Terrabacteria group>Firmicutes, A0A2U0RXR7 - Archaea>TACK group, A0A0F9ZY95 - Bacteria>unclassified, M7T7C2 and A0A1V4TQL4 - Archaea>Euryarchaeota both, B5YBS3 - Bacteria>Dictyoglomi. The shown region corresponds to the alignment part where heliorhodopsin 48C12 is fully represented, C and N termini of other heliorhodopsins are truncated. Helices for heliorhodopsin 48C12 are shown as coils, β-sheets as bold arrows, loops as plain lines.

**Fig. S13.**
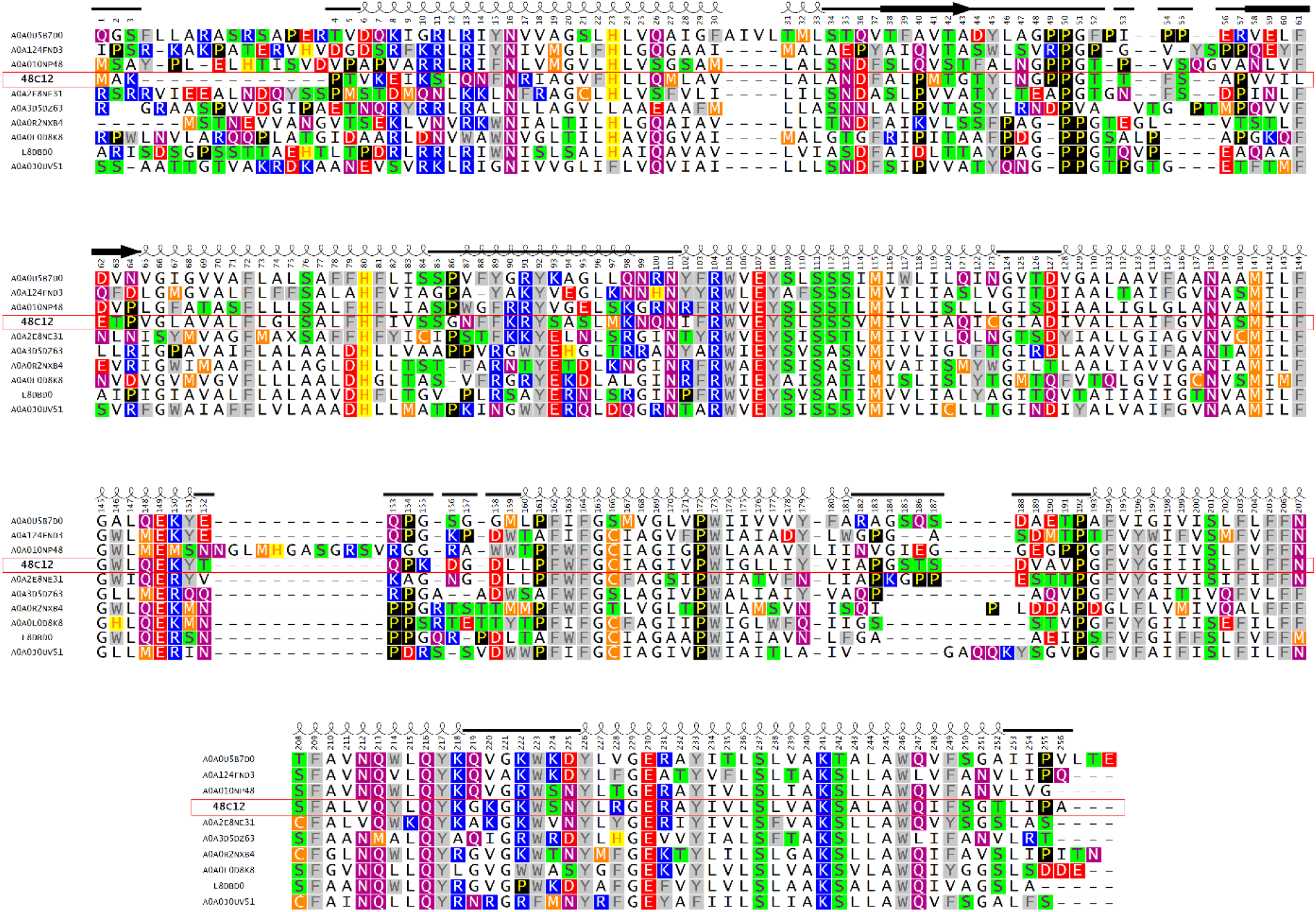
Sequence alignment of heliorhodopsin subfamily containing 48C12 heliorhodopsin (Heliorhodopsin 48C12 numbering). Proteins presented on the alignment are the most distant members of the subfamily 1. A0A0L0D8K8 – eukaryota, 48C12, A0A010NP48, A0A0J0UV51, A0A0R2NXB4, A0A0U5B7D0, A0A124FND3, A0A2E8NE31, A0A3D5DZ63 and L8DBD0-Bacteria>Terrabacteria group>Actinobacteria all. The shown region corresponds to the alignment part where heliorhodopsin 48C12 is fully represented, C and N termini of other heliorhodopsins are truncated. Helices for heliorhodopsin 48C12 are shown as coils, β-sheets as bold arrows, loops as plain lines.

**Fig. S14.**
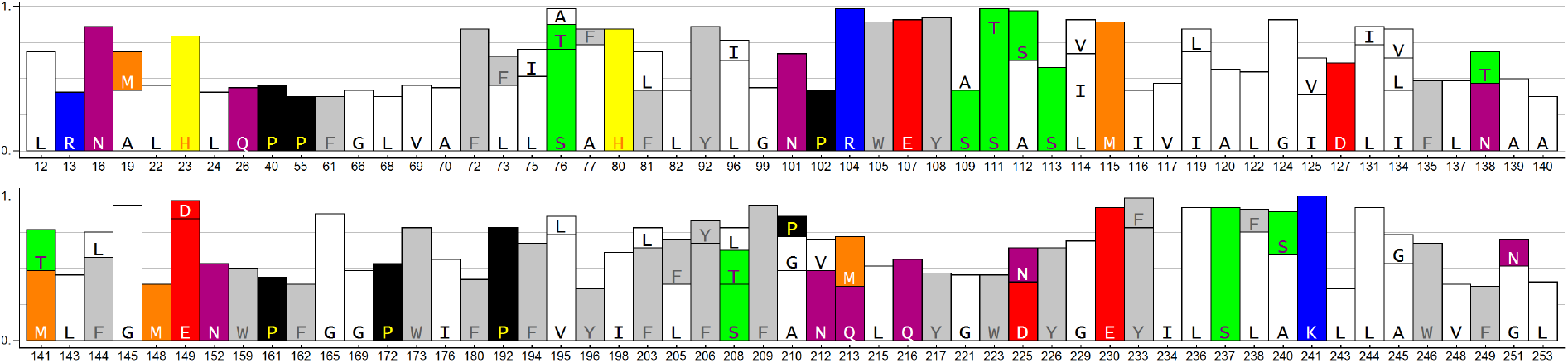
Most conservative residues in whole heliorhodopsins family. Whole base of heliorhodopsins (n = 479) was first processed to remove all sequences with more that 50% identity (n = 64) not to exaggerate conservativity of residues that are conservative among big subclasses of heliorhodopsins.

**Fig. S15.**
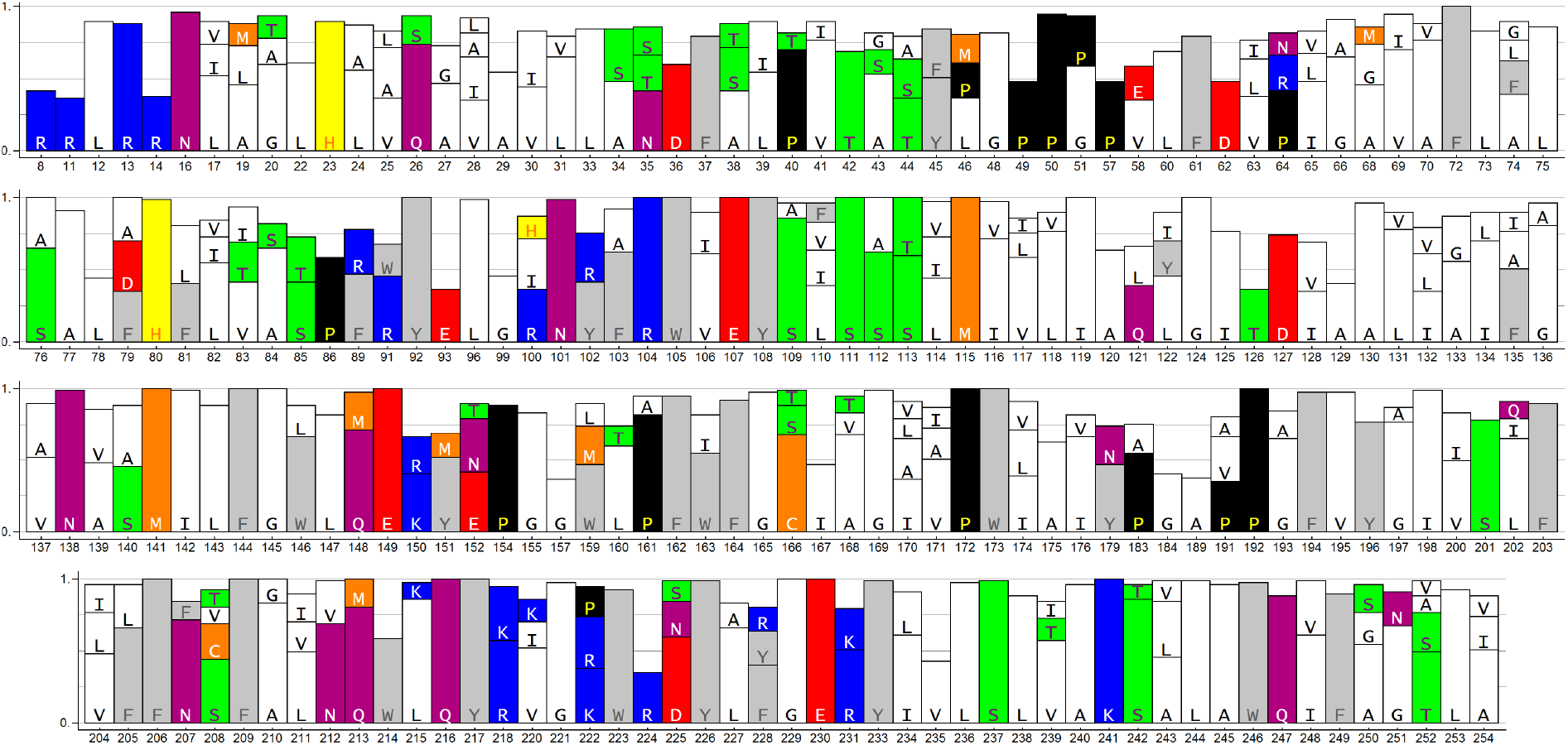
Most conservative residues in heliorhodopsins subfamily containing 48C12. Subfamily base (n = 195) was firstly processed to remove all sequences with more that 75% identity (n = 77) not to exaggerate conservativity of residues conservative in big subclasses of this subclass.

**Fig. S16.**
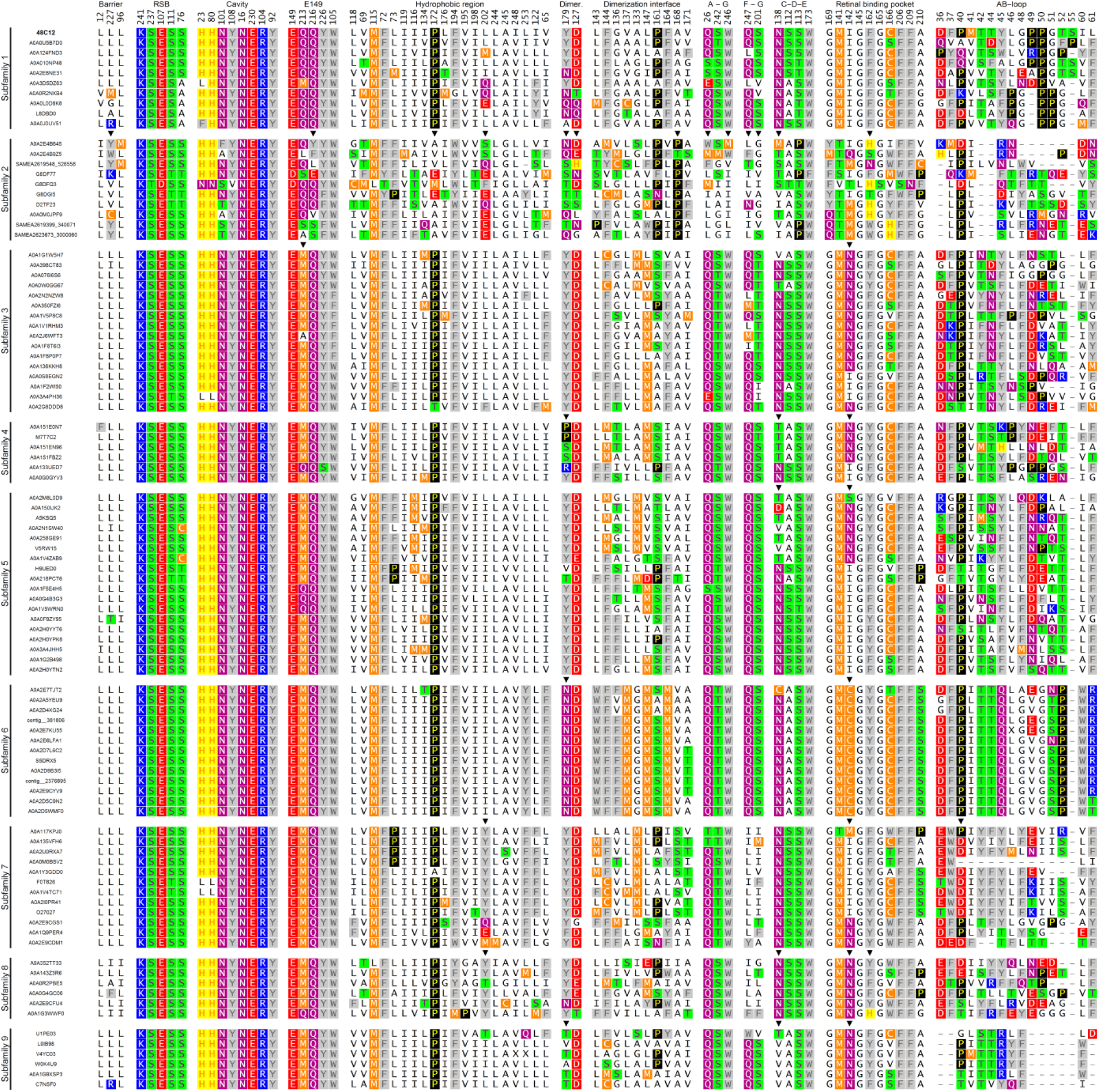
Alignment of HeRs from different classes. Shown positions chosen for structurally important regions: “Hydrophobic barrier between inner polar cavity and cytoplasm”, “Retinal and RSB-interacting residues”, “Residues, comprising inner polar cavity at the cytoplasmic part”, “Polar region at the cytoplasmic side near E149”, “Polar residues at the dimerization interface, responsible for contacts”, “Hydrophobic residues at the dimerization interface inside the membrane”, “Cluster between A and G helices at the extracellular side of the protein”, “Cluster between F and G helices at the extracellular side of the protein”, “Cluster near β-ionone ring of retinal between helices C, D, and E with 1 water molecule”, “Retinal binding pocket”, “AB-loop”

**Fig. S17.**
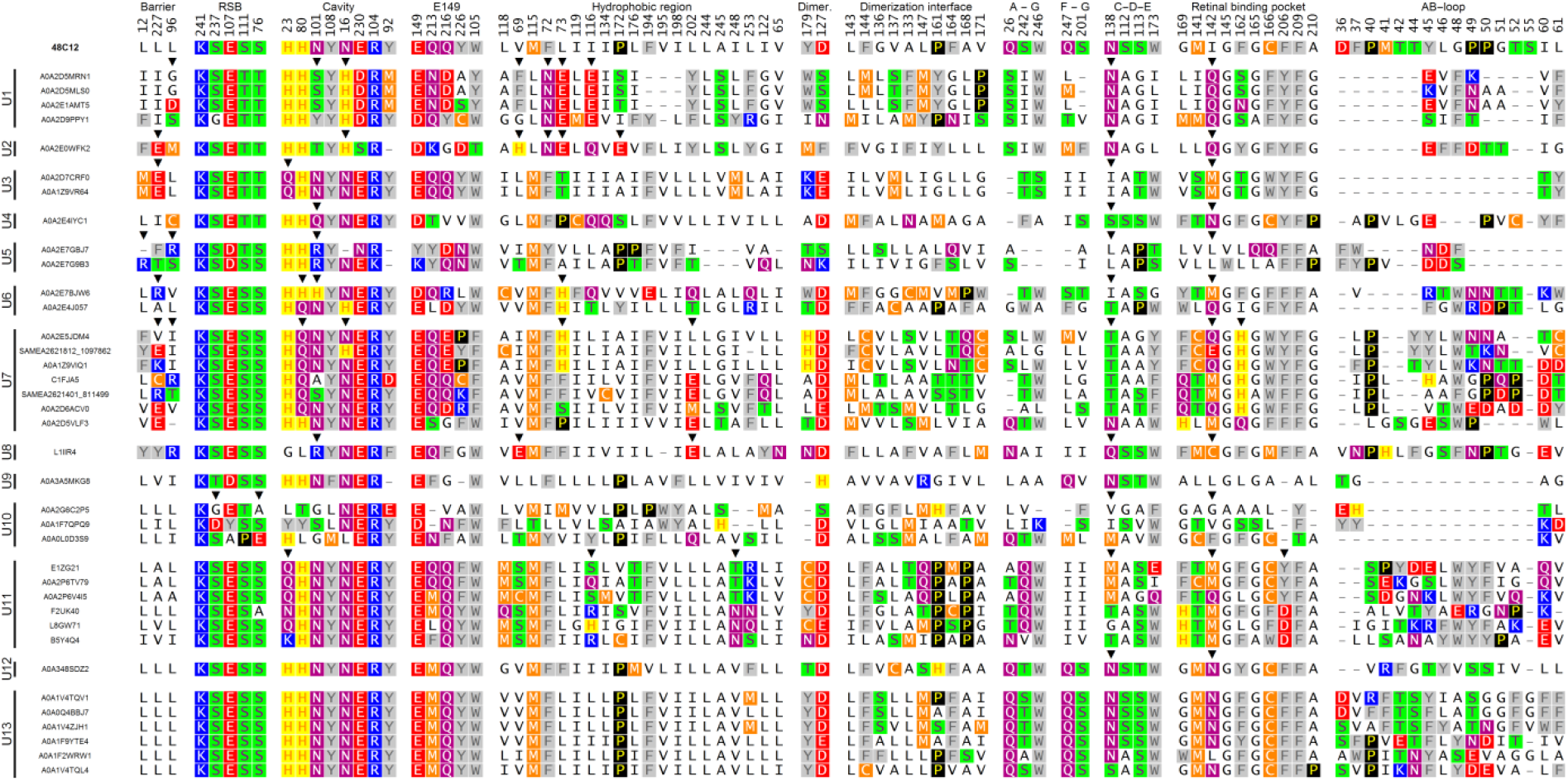
Alignment of HeRs from different classes from “Unsorted proteins”. Shown positions chosen for structurally important regions: “Hydrophobic barrier between inner polar cavity and cytoplasm”, “Retinal and RSB-interacting residues”, “Residues, comprising inner polar cavity at the cytoplasmic part”, “Polar region at the cytoplasmic side near E149”, “Polar residues at the dimerization interface, responsible for contacts”, “Hydrophobic residues at the dimerization interface inside the membrane”, “Cluster between A and G helices at the extracellular side of the protein”, “Cluster between F and G helices at the extracellular side of the protein”, “Cluster near β-ionone ring of retinal between helices C, D, and E with 1 water molecule”, “Retinal binding pocket”, “AB-loop”.

**Fig. S18.**
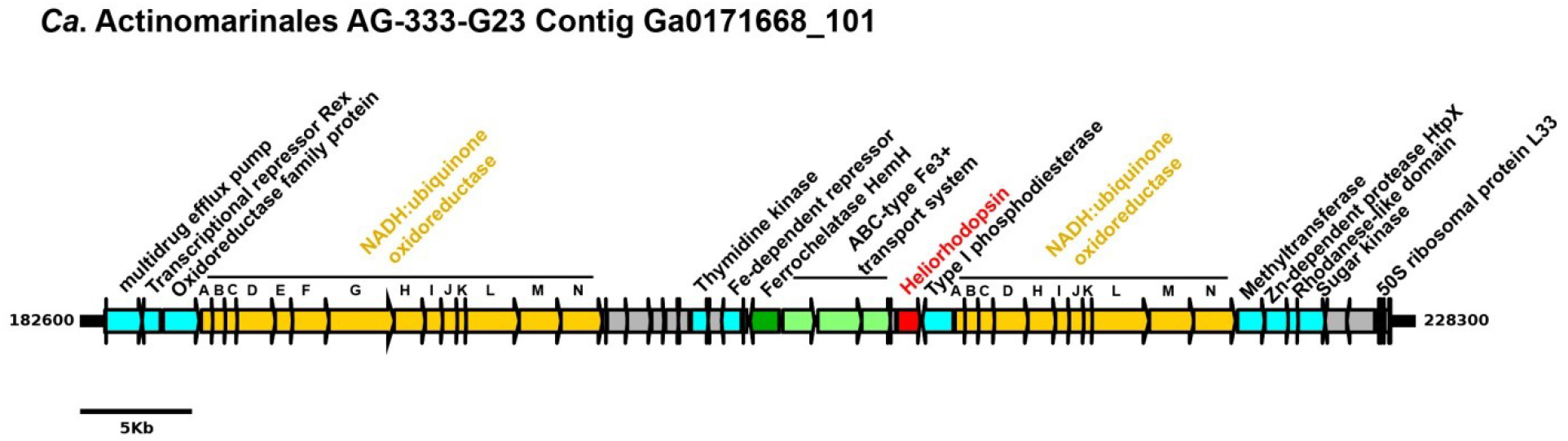
Linear representation of a genome fragment of 45.7 Kb retrieved from the marine *Ca.* Actinomarinales AG-333-G23 contig Ga0171668_101. Proteins were annotated against the NCBI-nr and InterPro databases. Genes coloured in grey represent hypothetical proteins. In red is coloured and highlighted the heliorhodopsin protein. A scale of 5 Kb long is represented in a black line below.

**Table S1.** Data collection and refinement statistics of the 48C12.

|  | *Violet form* | *Blue form* |
| --- | --- | --- |
| **Data collection** |  |  |
| Space group | P2<sub>1</sub> | P2<sub>1</sub> |
| Cell dimensions |  |  |
| *a*, *b*, *c* (Å) | 56.12, 59.75, 94.32 | 56.18, 60.74, 92.91 |
| $\alpha$, $\beta$, $\gamma$ (°) | 90, 92.30, 90 | 90, 92.02, 90 |
| Wavelength (Å) | 0.978 | 0.978 |
| Resolution (Å) | 49.06-1.50 (1.53-1.50) | 48.81-1.50 (1.53-1.50) |
| *R*<sub>merge</sub> (%) | 5.3 (113.8) | 7.0 (158.6) |
| *R*<sub>pim</sub> (%) | 3.4 (71.1) | 4.1 (93.3) |
| *I*/σ*I* | 8.3 (0.8) | 8.9 (0.7) |
| *CC*1/2 (%) | 99.9 (58.3) | 99.9 (33.3) |
| Completeness (%) | 98.6 (97.4) | 99.6 (95.0) |
| Multiplicity | 3.4 (3.5) | 3.8 (3.8) |
| Unique reflections | 98,436 (4795) | 99,767 (4713) |
| **Refinement** |  |  |
| Resolution (Å) | 20-1.50 | 20-1.50 |
| No. reflections | 93,455 | 94,710 |
| *R*<sub>work</sub>/*R*<sub>free</sub> (%) | 15.3/19.9 | 17.3/20.1 |
| No. atoms |  |  |
| Protein | 4066 | 4009 |
| Water | 233 | 162 |
| Lipid fragments | 409 | 420 |
| Retinal | 40 | 40 |
| Sulfate | 5 | 10 |
| Acetate | - | 8 |
| *B* factors (Å<sup>2</sup>) |  |  |
| Protein | 25 | 27 |
| Water | 36 | 35 |
| Lipid fragments | 36 | 53 |
| Retinal | 18 | 17 |
| Sulfate | 61 | 90 |
| Acetate | - | 29 |
| R.m.s deviations |  |  |
| Protein bond lengths (Å) | 0.005 | 0.004 |
| Protein bond angles (°) | 1.19 | 1.16 |
| Ramachandran analysis |  |  |
| Favored (%) | 98.0 | 97.5 |
| Outliers (%) | 0.4 | 0.4 |

